# Transcriptomics informs design of a planar human enterocyte culture system that reveals metformin enhances fatty acid export

**DOI:** 10.1101/2022.01.24.477515

**Authors:** Ismael Gomez-Martinez, R. Jarrett Bliton, Keith A. Breau, Michael J. Czerwinski, Ian A. Williamson, Jia Wen, John F. Rawls, Scott T. Magness

**Affiliations:** Joint Department of Biomedical Engineering, North Carolina State University and University of North Carolina at Chapel Hill 911 Oval Dr., Raleigh, NC, 27695 (USA); Department of Cell Biology & Physiology, University of North Carolina at Chapel Hill, Chapel Hill, NC, 27599 (USA); Department of Medicine, University of North Carolina at Chapel Hill, Chapel Hill, NC, 27599 (USA); UNC Center for Gastrointestinal Biology and Disease, University of North Carolina at Chapel Hill, Chapel Hill, NC, 27599 (USA); Department of Molecular Genetics and Microbiology, Duke Microbiome Center, Duke University School of Medicine, Durham, NC, 27710; Department of Biomedical Engineering, Woo Center for Big Data and Precision Health, Duke University Pratt School of Engineering, Durham, NC 27708

**Keywords:** Intestinal stem cells, absorptive enterocyte monolayers, fatty acid oxidation, drug screening

## Abstract

**Background & Aims:** Absorption, metabolism, and export of dietary lipids occurs in the small intestinal epithelium. Caco-2 and organoids have been used to study these processes but are limited in physiological relevance or preclude simultaneous apical and basal access. Here, we develop a high-throughput planar human absorptive enterocyte (AE) monolayer system for investigating lipid-handling, then evaluate the role of fatty acid oxidation (FAO) in fatty acid (FA) export, using etomoxir, C75, and anti-diabetic drug, metformin.

**Methods:** Single-cell RNA-sequencing (scRNAseq), transcriptomics, and lineage trajectory was performed on primary human jejunum. *In vivo* AE maturational states informed conditions used to differentiate human intestinal stem cells (ISCs) that mimic *in vivo* AE maturation. The system was scaled for high-throughput drug screening. Fatty acid oxidation (FAO) was pharmacologically modulated and BODIPY™ (B)-labelled FAs were used to evaluate FA-handling via fluorescence and thin layer chromatography (TLC).

**Results:** scRNAseq shows increasing expression of lipid-handling genes as AEs mature. Culture conditions promote ISC differentiation into confluent AE monolayers. FA-handling gene expression mimics *in vivo* maturational states. FAO inhibitor, etomoxir, decreased apical-to-basolateral export of medium-chain B-C12 and long-chain B-C16 FAs whereas CPT1 agonist, C75, and antidiabetic drug, metformin, increased apical-to-basolateral export. Short-chain B-C5 was unaffected by FAO inhibition and diffused through AEs.

**Conclusions:** Primary human ISCs in culture undergo programmed maturation. AE monolayers demonstrate *in vivo* maturational states and lipid-handling gene expression profiles. AEs create strong epithelial barriers in 96-Transwell format. FA export is proportional to FAO. Metformin enhances FAO and increases basolateral FA export, supporting an intestine-specific role.

## Introduction

The small intestinal (SI) epithelium is a selective barrier that serves as the point of entry for essential micro- and macronutrients to meet energy demands and preserve general homeostatic functions. Lipids are the most energy dense of the macronutrients and are absorbed by the SI epithelium^1^. The majority of dietary lipids are triglycerides (TAGs) and are broken down by the stomach and intestinal lumen into fatty acids (FAs) and monoglycerides^1^. FAs are then taken up by absorptive enterocytes (AEs), the predominant cell type of the SI epithelium^1^. Lipids can then be metabolized and used by AEs for cellular functions like energy production, membrane synthesis, and storage as lipid droplets^2^ or distributed to the body by the well-accepted lipoprotein-lymphatic system and by the less appreciated FA-portal vein pathway^3–5^.

Investigating uptake, metabolism, and export of dietary FAs, here collectively called ‘FA-handling’ *in vitro* is challenging due to the historical lack of physiologically relevant culture models. As metabolic disorders such as dyslipidemia, diabetes and obesity^1,6,7^ are on the rise^8,9^, there is strong interest in evaluating how genetics^10–12^ and environmental factors, such as alterations in gut microbiota^13–16^ and eating behaviors^1,6,7^, are associated with FA-handling mechanisms, which by nature are complex to study in humans or animal models. Limitations of 3D organoid cultures, ethical considerations of human research, and inadequacies of animal models compound the challenges and limit scientific progress toward solutions for these and other metabolic diseases. In this regard, an AE cell culture platform using primary human cells and coupled with simple and sensitive detection of FAs and their metabolites would represent a significant improvement and address many of the limitations of existing *in vitro* culture models.

Traditionally, cell culture models of human SI epithelium have largely relied on cancer or immortalized murine cell lines (*i.e.,* Caco-2^17^, ICCL-2^18^, etc.), which retain properties consistent with undifferentiated states^19,20,21^. Organoid culture models have become popular alternatives because they are typically derived from primary intestinal epithelial stem cells (ISCs) and can differentiate into the main mature lineages of the differentiated gut epithelium^22,23^. Organoids are small (~100-1000μm diameter) spherical structures cultured in thick hydrogels. They are comprised of polarized cells where the enclosed apical or ‘luminal’ surface precludes application of FAs to the physiologically relevant surface to mimic dietary absorption^23^. Organoids can be cultured with the apical surface facing outward (apical-out organoids)^24^; however, this causes the basal surface to be enclosed and inaccessible, preventing sampling of exported or metabolized lipids. Collectively these factors limit interpretations, reduce throughput, and prohibit analyses necessary to accurately assess FA-handling across the AE monolayer.

Conventional methods to detect FAs has relied on using radioisotope- and heavy isotope-labeled FA analogs^25–31^. These isotope-labeled FAs are thought to behave similarly to native FAs in absorptive, metabolic, and export processes^25–31^; however, special safety precautions and sophisticated downstream analytics (*e.g.,* mass spectrophotometry) are required, limiting access of these assays to many investigators. Fluorescently-labeled FAs represent an attractive alternative because they do not require special handling, are sensitive, commercially available, and can be detected using a variety of common instruments and methods (*e.g.,* plate readers, microscopes, thin layer chromatography)^32^. For example, BODIPY™ (B) is a brightly fluorescent fluorophore, and B-FA analogs have been shown to mimic endogenous FA metabolism and transport, making it an effective tracer for FAs in lipid-handling studies^32–36^. Importantly, unlike isotopically labeled FAs, fluorescently labeled FAs also permit imaging of FA-handling in live and fixed cells^35,36^.

Our group has recently developed methods for culturing and indefinitely expanding primary human ISCs as 2D monolayers^37^. ISCs cultured this way can then be transferred to Transwell™ inserts, cultured to confluence, and then terminally differentiated into absorptive and secretory lineages found *in vivo*^37,38^. Monolayers cultured on permeable Transwell inserts are in contact with apical and basal reservoirs where factors, drugs, and metabolites can be easily added or sampled throughout an experiment^37^. Unlike organoids, monolayers grow as planar sheets rather than spheres suspended in thick hydrogels, allowing use of imaging systems that are in common use in basic science laboratories, robotic drug screening and validation platforms.

Our study here has two primary goals; the first is to develop and validate a new culture system to study FA-handling, and second is to use the system to demonstrate utility for evaluating the impact that a set of drugs has on FA oxidation (FAO) and export of FA metabolites. We take an approach that first defines the baseline transcriptomic state of relevant lipid-handling genes and then tailor culture conditions to mimic the *in vivo* lipid-handling gene profiles. Readouts for FA-handling are designed for practicality, sensitivity, and high-throughput applications. Using this new system, we pharmacologically inhibit and potentiate FAO and observe changes in FA export that informs a hypothesis that FAO increases export of medium- and long-chain FA metabolites across the basolateral membrane. This is tested and the findings reveal new biological insights into the role of FAO on export of FAs with implications for understanding blood-glucose regulation and appetite control.

## Results

### Single-cell transcriptomics of jejunal and ileal human mucosa define early, intermediate, and mature nutrient-handling enterocytes

First, we sought to characterize the baseline transcriptomic profiles of human AEs *in vivo* to guide the development of an *in vitro* model of human FA-handling. The distal SI (jejunum and ileum) represents the majority of absorptive epithelium in the human SI^39^; however, lipid-handling transcriptional profiles of jejunal and ileal absorptive lineages have not been fully described at the single-cell level. Endoscopic biopsies have enabled single-cell transcriptomics to be performed on human duodenal^40^ and ileal^41^ mucosa; however, single-cell transcriptomics has only recently been reported for the jejunum^42,43^, with limited characterization of absorptive function^39,43.^. To further define absorptive function of the distal SI, single-cell RNA-sequencing (scRNAseq) was performed on primary jejunal and ileal epithelium isolated from a healthy organ donor (Fig. S1). Annexin V staining demonstrated high viability (95.7%, Fig. S1) of single cells (Fig. S1A-E) with 1,788 having passed quality filtering (Fig. S1F, G). To identify the distinct cell types captured in the dataset, dimensional reduction was performed^44^, revealing that the cell populations represented the major reported cell lineages in the human SI epithelium^40^ (Fig. 1A). Most cells in the dataset clustered separately from the minority secretory lineages (enteroendocrine, goblet, BEST4+, and tuft). This main cluster was comprised of cells consistent with ISCs, transit-amplifying (TA) cells, and AEs (Fig. 1A). The high viability, quality and capture of the full complement of major SI epithelial lineages provided a strong foundation for transcriptomic characterization.

**Figure 1.**
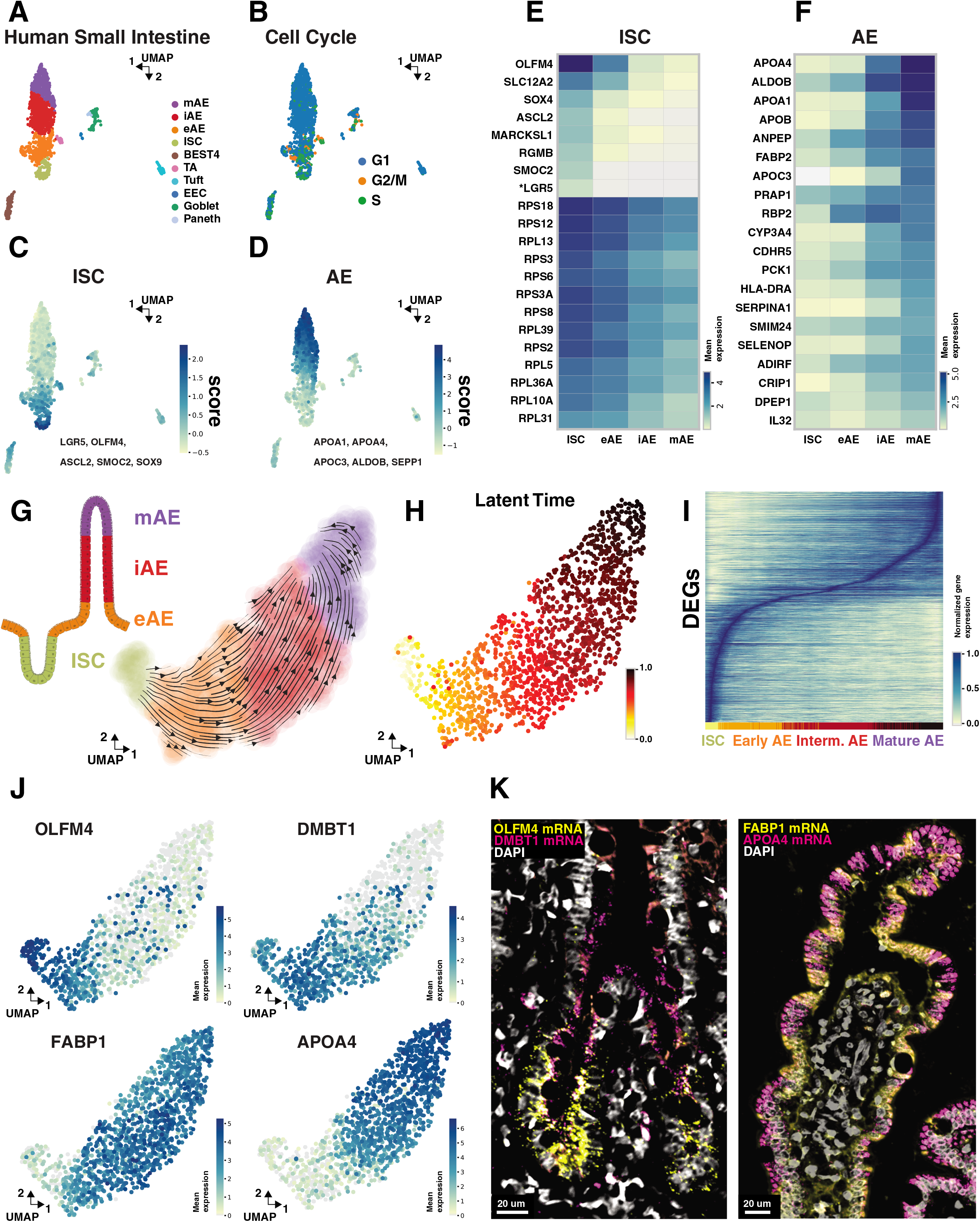
Single-cell transcriptomics of healthy human absorptive epithelium. **(A-D)** Unbiased Leiden clustering of primary jejunum and ileum. **(A)** Identification of cell types. **(B)** Inferred cell cycle state based on expression of previously categorized G1, G2M, and S phase associated genes^46^. **(C)** ISC score based on expression of established ISC genes *LGR5, OLFM4, ASCL2, SMOC2, SOX9*. **(D)** AE score based on expression of established AE genes *APOA1, APOA4, APOC3, ALDOB, SEPP1*. **(E-F)** Gene expression of ISC, and AE populations identified in 1A. **(E)** Expression of top 20 ISC genes previously identified in human ileum^41^. \**LGR5* was not within the top 20 ISC genes previously identified in human ileum, but it was included for reference. **(F)** Expression of top 20 AE genes previously identified in human ileum^41^. **(G)** Bottom right, vectors calculated based on a dynamical model of RNA velocity showing likely cell transitions of ISCs, eAEs, iAEs and mAEs. Top left, schematic showing likely positions of identified ISC and AE populations along the crypt-villus axis. Note that secretory and ileal lineages were removed from the analysis to focus on the relationship between ISCs and maturing of jejunal AEs. **(H)** A latent time value between 0 and 1 was assigned to each cell to order cells along the trajectory modeled by RNA velocity. A latent time of 0 means that the cell has yet to experience any differentiation in the modeled trajectory whereas a latent time of 1 means that the cell has progressed completely through the modeled differentiation pathway. **(I)** Identified human jejunal AEs were ordered based on latent time (x-axis) and DEGs of identified human ISC and AE populations were plotted on the y-axis. *LGR5 was not among the top 20 ISC genes identified in human Ileum^41^ **(J)** UMAPs showing expression of select DEGs from population of cells along the ISC to mAE differentiation axis. **(K)** *In situ* hybridization of ISC and eAE DEGs *OLFM4* and *DMBT1* respectively (left). *In situ* hybridization of iAE and mAE DEGs *FABP1* and *APOA4* respectively (right).

In human duodenum and ileum, AEs are sub-categorized as early (eAE), intermediate (iAE), and mature (mAE) based on maturation state^40,45^. Genes associated with these three AE sub-sets were also observed in jejunal and ileal Leiden-clustering (Fig. 1A). Jejunal and ileal AEs clustered together demonstrating a high degree of transcriptomic similarity between these cells (Fig. S1H). Since Leiden-cluster boundaries appear artificially binary, each cell was independently interrogated using a computational score comprised of curated gene sets that would predict cell lineages for each Leiden-cluster. Three different curated gene sets from prior studies were used to identify different cell cycle stages^46^, ISCs^47–49^, and AEs^41^ (Fig. 1B-D). The cell-cycle score showed most cycling cells are strongly associated at one end of the cluster, which is consistent with cells exhibiting the strongest ISC score (Fig. 1B,C). By contrast, cells with the highest AE score were strongly associated with the opposite end of the same cluster (Fig. 1D). Twenty ISC signature genes that were reported to be the most highly expressed genes in human ileal ISCs^41^ were also highest in jejunal ISCs (Fig. 1E). These ISC-associated genes gradually decreased in jejunal AE populations consistent with differentiation (Fig. 1E). Conversely, 20 of the most highly expressed AE signature genes identified in human ileal mAEs^41^ were lowest in jejunal ISCs. These AE-associated genes gradually increased in jejunal AE populations as they differentiated. (Fig. 1F). Together these data suggest jejunal AEs follow a similar maturation trajectory as duodenal and ileal cells from other reports^40,41,45^.

### Computational lineage trajectory analyses reveal human absorptive enterocytes progress through maturation stages with increasing FA-handling gene expression

A recent study of murine jejunal villi described an AE maturation program wherein spatially distinct zones along the villus were used to define AE functions (*e.g.*, antimicrobial functions towards the base, lipoprotein secretion towards the tip)^50^. Whether human AEs also progress through maturation states corresponding to distinct functions along the length of the villus is unknown. Progressive expression of mature AE markers in our jejunal dataset (Fig. 1F) led to the hypothesis that human AE clusters correspond to lineages with distinct gene expression profiles of maturation along the villus. To test this hypothesis, we used a computational framework based on the rate of change of spliced and unspliced mRNA ratios (*i.e.,* RNA velocity)^51^ and a differentiation-specific RNA velocity-based metric (*i.e.,* latent time) (Fig. 1G,H)^52^. Human jejunal cells were used in the analysis to be consistent with previous mouse studies^50^. Vectors calculated based on the solved dynamical model of RNA velocity predict a trajectory of ISCs gradually differentiating to mAEs (arrows, Fig. 1G). Next, a latent time value between 0 and 1 was assigned to each cell to order cells along the trajectory modeled by the RNA-velocity vectors (Fig. 1H). A latent time of 0 means that the cell has yet to enter the modeled trajectory whereas a latent time of 1 means that the cell has progressed completely through the modeled differentiation pathway. Through combining RNA velocity and latent time, we demonstrate a single path of likely cell transitions from ISCs into eAEs, followed by iAEs and culminating with mAEs (Fig. 1G,H).

To visualize the number of differentially expressed genes (DEGs) between these maturation states (*i.e.,* eAE, iAE, and mAE), the 1,537 identified DEGs from these populations were plotted against all jejunal ISCs, eAEs, iAEs and mAEs as ordered by the calculated latent time (Fig. 1I, Fig. S2A-D, Table S3). ISCs and mAEs had the largest amount of DEGs (732 and 592, respectively) with eAEs and iAEs only having 106 and 107 DEGs, respectively (Table S3). The pattern of gene expression showed that genes highest in ISCs gradually turn off as the cells mature (Fig. 1I). Conversely mature AE genes begin to turn on in the eAE and iAE states (Fig. 1I). The cell maturation analyses were next further refined using curated gene sets specific for FA handling (Fig. S3A). Expression of genes associated with chylomicron assembly (APOA1, APOA4, APOB) were enriched in iAE and mAE populations, while distinct subsets of genes involved in regulating FA transport (CD36, SLC27A1, SLC27A5) and FAO (CPT1A, PPARG, PPARGC1A) were differentially enriched in each maturation state, suggesting discrete regulation of these lipid-handling mechanisms along the villi (Fig. S3A).

*In situ* hybridization was performed on jejunal tissue sections from the same donor to validate the predicted computational trajectory by locating marker genes for each maturation stage to points along the villus (Fig. 1J,K). Consistent with our predictions, *OLFM4 (ISC marker)* localized to the crypt base, *DMBT1* (antimicrobial function) was found in the upper crypt and lower villus, *FABP1* (lipid chaperone) was enriched in the mid-villus region, and *APOA4* (chylomicron assembly) was enriched at the villus tip (Fig. 1K). Together these findings support distinct transcriptional states and associated lipid-handling mechanisms with each AE maturation stage along the crypt-villus axis in the human jejunum. Consistent with findings in mice, human AEs appear to perform distinct functions as they differentiate along the crypt-villus axis^50^.

### Characterization of culture model by scRNAseq demonstrates differentiation of human jejunal ISCs generates highly pure monolayers of absorptive enterocytes

In prior work, our group developed platforms that promote robust, long-term expansion of human ISCs^37^. When cultured on Transwell culture systems, ISCs can be induced to differentiate into AEs with strong barrier function and used for transport studies^53,38^ (Table S1). The Transwell membrane allows for application and retrieval of FAs and their metabolites from the apical and basal reservoirs, respectively. Both are crucial parameters not available in organoids that enable mechanistic studies of FA-handling *in vitro*.

To define cell phenotypes and characterize lineage purity at single-cell resolution, scRNAseq was performed on ISCs under expansion conditions^53^ (Table S2) and AE monolayers after 5-days of differentiation (Fig. 2A). Inhibition of Notch signaling is required for specification of the secretory lineage^54,55^. Because we initiated differentiation by removal of ISC growth factors without addition of a Notch inhibitor, we hypothesized that ISCs would follow an absorptive rather than secretory differentiation trajectory. Leiden clustering of ISC expansion and differentiation conditions showed two distinct clusters unique to each media formulation (Fig. 2B). 40% of cells in the ISC cluster were predicted by computational methods to be in either S- or G2/M-phase, whereas >99% of cells in AE differentiation conditions were predicted to be in G0/G1-phase (Fig. 2C).

**Figure 2.**
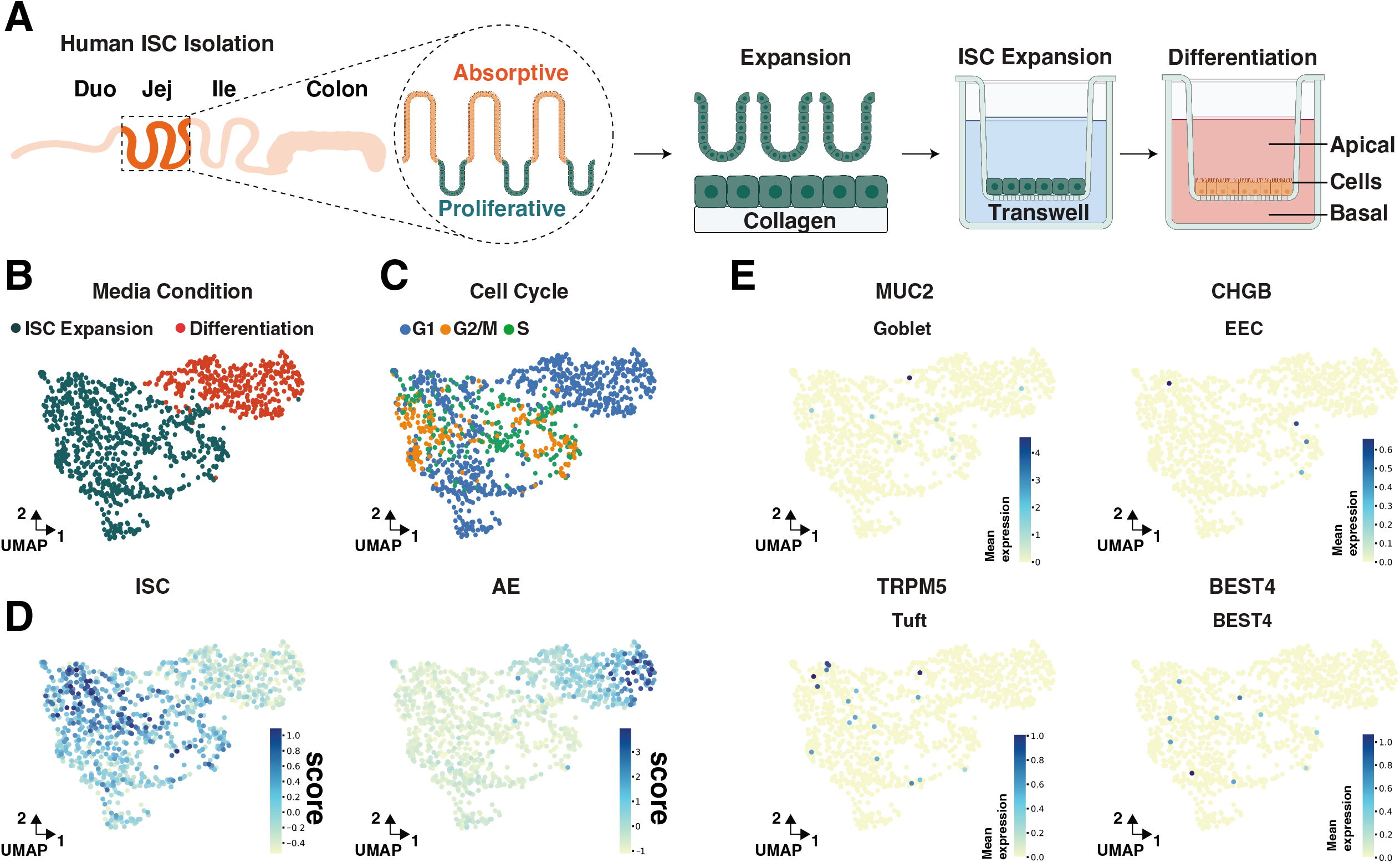
Single-cell transcriptomics of 2D ISC and AE cultures. **(A)** Schematic of isolation, expansion, and differentiation of ISCs from the jejunum of a healthy organ donor. **(B)** Unbiased Leiden clustering of collagen grown ISCs (green) and ISCs differentiated for 2 days (orange). **(C)** Inferred cell cycle state based on expression of previously categorized G1, G2M, and S phase associated genes^46^. **(D)** Left, ISC score based on expression of established ISC genes *LGR5, OLFM4, ASCL2, SMOC2, SOX9*. Right, AE score based on expression of established AE genes *APOA1, APOA4, APOC3, ALDOB, SEPP1*. **(E)** Expression of secretory lineage markers *MUC2, CHGB, TRPM5, BEST4*.

Computational cell cycle predictions cannot distinguish between cells in G1 or cells that have left the cell cycle, thus, scoring of ISC- and AE-gene profiles was performed to assess if differentiation conditions were sufficient to confer terminal differentiation towards an absorptive fate (Fig. 2D). ISC and AE scores were generated for cells grown in *in vitro* monolayers using curated gene sets that identify either ISCs or AEs (Fig. 2D). This scoring revealed that *in vitro* monolayers showed higher expression of ISC genes in expansion conditions and higher expression of AE genes in differentiation conditions (Fig. 2D). Very rare cells expressing markers of classic secretory lineages (goblet cells, enteroendocrine cells, tuft cells, BEST4^+^ cells) were observed (Fig. 2E). These data demonstrated that culture conditions promoted ISC differentiation towards an absorptive cell fate and indicate the potential to model physiological AE differentiation *in vitro*.

### Culture methods developed in 96-Transwell format drive time-dependent ISC differentiation and AE maturation described by transcriptomic states

AE planar monolayers were scaled to a 96-well Transwell format for high-throughput applications. Transepithelial Electrical Resistance (TEER) was used to monitor barrier integrity^56^ over 15 days of ISC expansion (4 days) and AE differentiation (12 days) (Fig. 3A). As expected^57^, ISC conditions produced a stable and low TEER, while differentiation media (DM) produced an immediate and progressive increase in TEER that peaked at 7 days of differentiation (Fig. 3A). The progressive increase in TEER suggests changes in gene expression patterns consistent with mature AEs. To confirm this at the transcriptomic level, bulk RNA sequencing (bulk RNAseq) was performed on AEs in differentiation conditions for 0, 2, 5, 7, 10, and 11 days (Fig. 3B-D). Principal component analysis (PCA) demonstrated tight agreement between technical replicates and showed rapid and large transcriptomic changes following just 2 days of differentiation (Fig. 3B). Bulk RNAseq of AE monolayers revealed a trajectory of progressive transcriptional changes through time (Fig. 3B), suggesting an intrinsic program of progressive AE maturation that begins upon removal of ISC niche growth factors.

**Figure 3.**
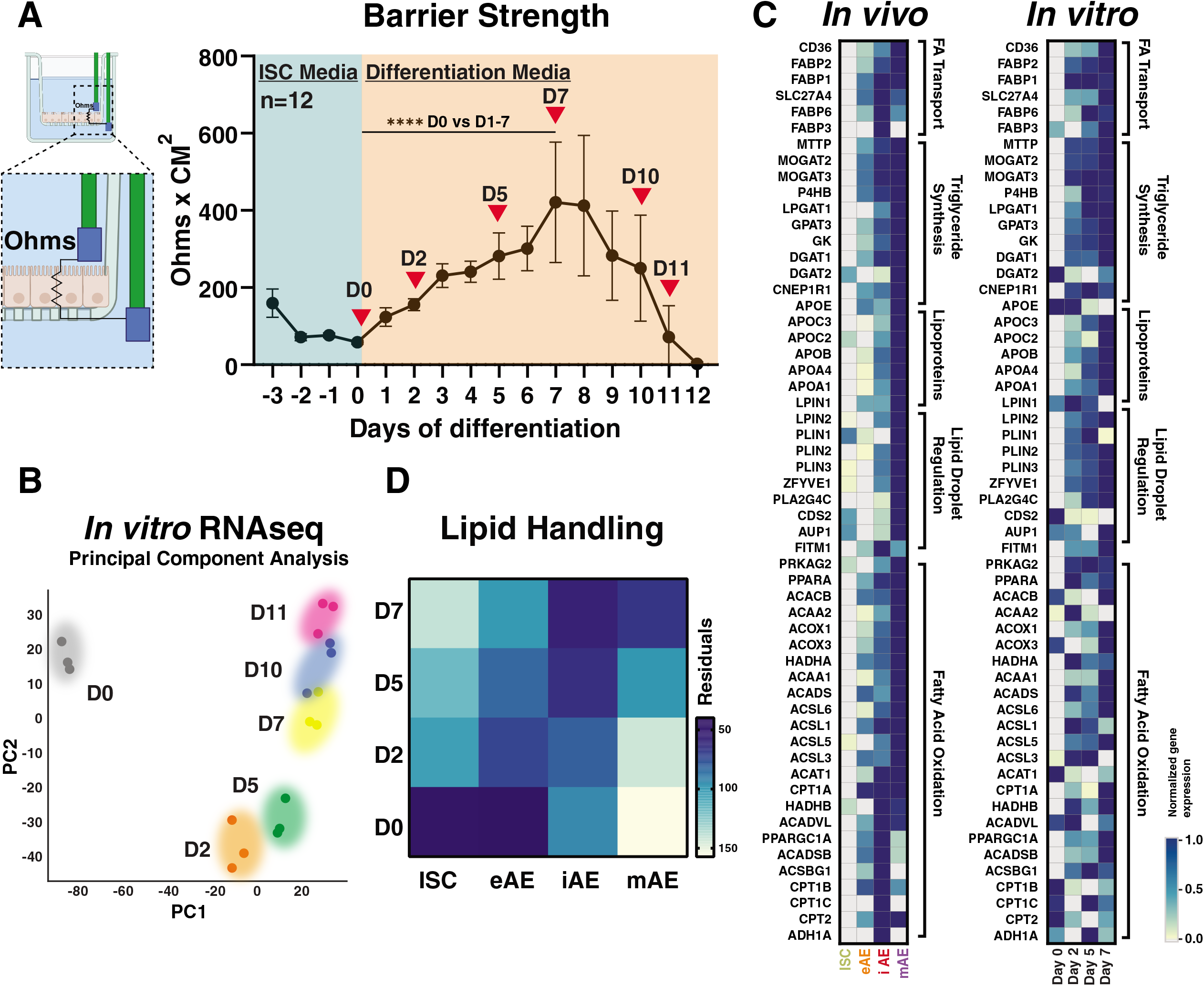
RNA-sequencing reveals a time dependent AE maturation program in jejunal ISCs *in vitro*. **(A)** Transepithelial electrical resistance (TEER) of jejunal ISCs n=12 Transwell inserts. ISCs were maintained in expansion media for 3 days then switched to differentiation media for 12 days. TEER was monitored daily. Red triangles indicate time points where cells were lysed for bulk RNA-seq. A Kolmogorov-Smirnov test was performed comparing TEER between D0 and D1-D7. **** p < 0.0001. **(B)** Principal component analysis of sequenced transcriptomes. ISCs (grey) differentiated for 0 (D0), 2 (D2), 5 (D5), 7 (D7), 10 (D10), and 11 (D11) days. (C) Expression of FA-handling genes *in vivo* (left) and *in vitro* (right). **(D)** Residual plot of key lipid-handling genes. Linear correlation was used to quantitatively describe the extent of similarity between FA-handling genes expressed during *in vivo* and *in vitro* AE maturation.

Next, key FA-handling gene expression profiles were compared between *in vivo* scRNAseq data and *in vitro* bulk RNAseq data to further inform optimal time points to investigate FA-handling. Genes associated with pathways of FA-handling (*i.e.,* FA transmembrane transport, FAO, lipid droplet formation, chylomicron secretion and TAG metabolic processes) generally increased in expression over time both *in vivo* and *in vitro* (Fig. 3C, D). There appeared to be a strong correlation in the magnitude of gene expression between mAEs *in vivo* and D7 AEs *in vitro* (Fig. 3C). Linear correlation was used to quantitatively describe the extent of similarity between FA-handling genes expressed during *in vivo* and *in vitro* AE maturation (Fig. S3B). Pairwise comparisons of mean lipid-handling gene expression values were made for each *in vivo* differentiation state and each *in vitro* differentiation state (Fig. S3B) revealing that D7 AEs are more similar to intermediate- and mature-AEs than ISCs and eAEs, providing evidence that *in vitro* directed differentiation mimics *in vivo* AE differentiation with respect to lipid-handling processes. (Fig. 3D). Together, these data show that 7-days of *in vitro* differentiation is sufficient to establish epithelial monolayers and lipid-handling gene expression patterns similar to *in vivo* AEs.

### High-throughput planar AE cultures produce robust epithelial barriers and preserve FA-handling functions detected by fluorescent FA conjugates

High TEER and transcriptomic data suggest that AE monolayers could serve as an effective *in vitro* model for human AE lipid-handling. Strong barrier function is required for accurate interpretation of FA-handling by AEs, since a leaky barrier would allow FAs to bypass the AE monolayer. As a means to provide sensitive readouts for FA-handling that is accessible to most laboratories, we adopted fluorescent analogs that can be easily detected by fluorescent plate reader and thin layer chromatography. To validate barrier function, 10kD Dextran conjugated to Alexa Fluor™ 647 (Dextran-647), a non-absorbable fluorescent polysaccharide, was applied to the apical monolayer surface followed by quantification of basolateral fluorescence at 2, 4 and 6 hours of apical exposure (Fig. 4A). There were nearly undetectable levels of Dextran-647 (< 0.2 pmols) in the basal reservoir at any time point, whereas Transwells without cells allowed > 30 pmols to diffuse through and reached an equilibrated concentration within ~4 hours. These data demonstrate strong barrier function and show that significant epithelial barrier defects can be readily detected by Dextran-647 signal in the basal reservoir as early as 2-hours post-exposure.

**Figure 4.**
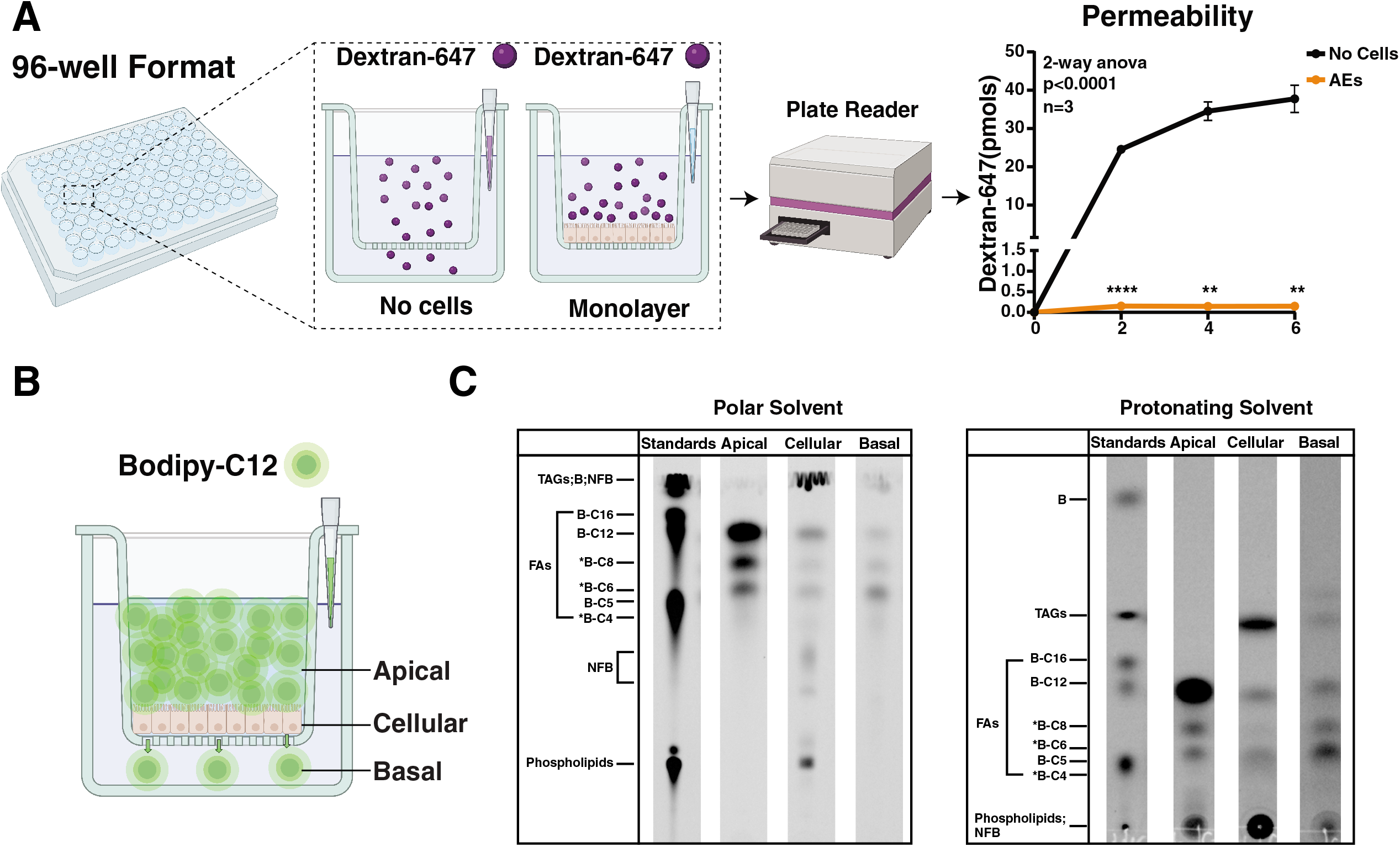
Application of fluorescent polysaccharide and FA reveal integral barrier and FA-handling *in vitro*. **(A)** Left, schematic showing application of Dextran-647 to the apical surface of AEs or empty Transwell cultures. Right, subsequent sample collection and fluorescence quantification via plate reader. ISCs were differentiated for 7 days before application of 1uM Dextran-647. 50uls of media from the basal reservoir were removed at 2, 4 and 6 hours from empty Transwell insert s or AE cultures and replaced with fresh differentiation media; ** p < 0.005, **** p < 0.0001. **(B)** Schematic of B-C12 application to AE Transwell cultures and subsequent retrieval from apical, cellular, and basal reservoirs. **(C)** Thin layer chromatography of apical, cellular and basal reservoirs of AE Transwell cultures after 6 hours of B-C12 application using polar and protonating solvents. A previous study^62^ using TLC to identify B-lipid species with the same solvents used in this study reported naturally fluorescent bands that do not correspond to B-lipids used and are labelled here as NFB. *Location of B-C4, B-C6 and B-C8 were inferred from the experiment in supplemental figure 4. Experiments were performed in triplicate (n = 3 AE transwell cultures).

BODIPY (B) is a bright fluorophore that has a different fluorescent signature than Dextran-647, facilitating separate detection of these two molecules by plate readers. B-FA analogs have been used extensively for lipid-trafficking and metabolism studies *in vitro* and *in vivo* and are considered to be accurately handled by absorption, metabolism, and export mechanisms^58–62^. Thus, to characterize FA-handling properties of AE monolayers, we applied the MCFA analog B-C12 to the apical surface of AE cultures for 6 hours (Fig. 4B). Apical (input) and basal (output) media was collected, and AEs were lysed after 6 hours of apical exposure (Fig. 4B). Thin layer chromatography (TLC) was used detect and quantify input B-FAs as well as metabolized and exported B-FA products. A combination of polar and protonating solvents were used to resolve the complex mixture of B-FAs and metabolites (Fig. 4C). B-labeled FA standards were applied to the TLC plate in a separate lane and unknown FA-species of different carbon-chain lengths were extrapolated (Fig. 4C, S4). These methods were scaled for the 96-well Transwell system using less than 170 μl of media or cell lysate. A previous study^62^ using TLC to identify B-lipid species with the same solvents used in this study reported naturally fluorescent bands that do not correspond to B-lipids and are labelled here as NFB (Fig. 4C). TLC demonstrated clear separation of key FA-species in all three reservoirs (*i.e.,* Apical, Cellular, Basal) indicating robust sensitivity of this approach for detection of a broad range of FA-species (Fig. 4C).

To evaluate FA-handling by AE monolayers, B-C12 was incubated with AE monolayers for 6 hours followed by TLC analysis to identify the B-FA or B-metabolites in each reservoir (*i.e.,* Apical, Cellular, Basal). TLC demonstrated that most of the B-labeled species in the apical reservoir were medium-chain FAs (B-C12, B-C8 and B-C6) (Fig. 4C). A short-chain species consistent with B-C4 was also detected in the apical reservoir at a lower level (Fig. 4C). In the intracellular reservoir, the largest species were TAGs, indicating robust FA esterification, and phospholipids indicative of B-C12 being metabolized and incorporated into the lipid bilayers^63^. In the cell lysates, FAs were some of the lowest B-lipid species suggesting dynamic metabolism, diffusion, and mobilization. In the basal reservoir, FAs were the predominant B-lipid species with a smaller fraction consisting of TAGs suggesting basal export of FAs and chylomicrons (Fig. 4C). Together these data demonstrate robust detection of input B-FAs, a broad range of the derivative metabolites, and their relative distributions in each reservoir as the B-lipids are processed by AEs.

### Inhibiting FAO in AE monolayers decreases basolateral export of oxidized FA species

We next explored the utility of the platform for investigating small molecule perturbations on FAO. Etomoxir, a CPT1 (Carnitine palmitoyl transferase 1) inhibitor, was used to inhibit FAO. When CPT1 is inhibited, FAs cannot be imported into the mitochondria where FAO normally catabolizes longer-chain FAs to smaller-chain FAs^64^ (Fig. 5G). *CPT1A* is robustly expressed in primary human AEs both *in vivo* and *in vitro* at Day 7 of differentiation (Fig. 3C). After pretreatment with etomoxir, a variety of different chain-length B-FAs (SCFA; B-C5, MCFA; B-C12 or LCFA; B-C16) were applied to the apical reservoir to mimic postprandial AE exposure to FAs *in vivo*. Media from the basal reservoir was taken at 2-, 4- and 6-hours after application of fluorescent FAs. The impact of etomoxir on basal export of FAs and FA metabolites was evaluated in real-time by total fluorescence detection by plate reader, and TLC was used to identify Bodipy-labeled metabolites using small volumes of media from the 96-well Transwell platform (Fig. 5A).

**Figure 5.**
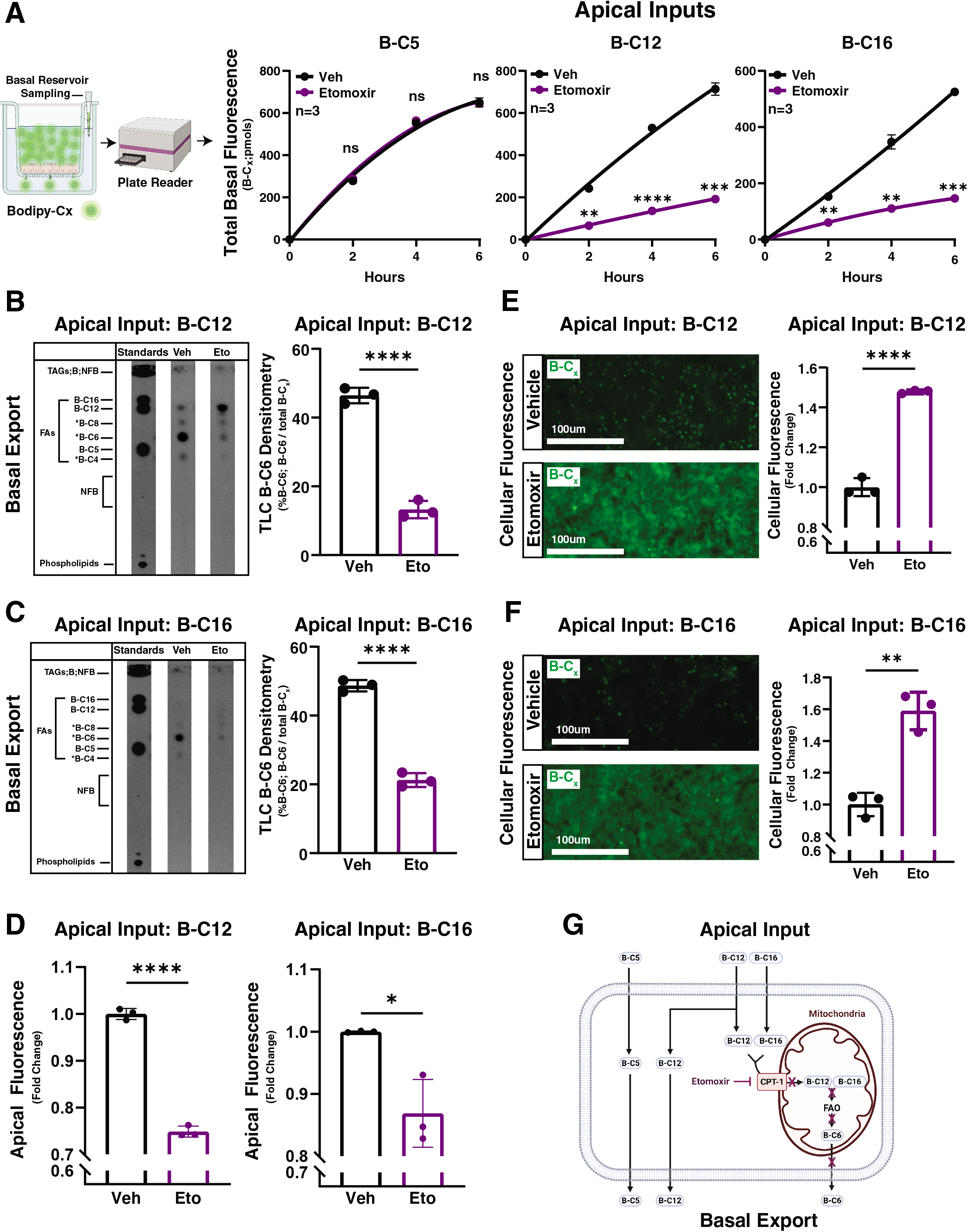
Inhibiting FAO reduces FA export. **(A)** Left, schematic showing basal reservoir sampling following application of B-FAs to the apical surface of AE Transwell cultures and subsequent quantification of basal fluorescence via plate reader. Right, quantification of basal fluorescence from B-C5, −C12, and −C16 treated AE cultures treated with vehicle (DMSO), or etomoxir (Eto). ** p < 0.01, *** p < 0.005, **** p < 0.0001. 50ul of basal media were collected and quantified by plate reader at 2-, 4- and 6-hours post FA application. **(B, C)** Left, TLC of basal-reservoir media collected 6 hours post vehicle or etomoxir and B-C12 or B-C16 application. Right, quantification of the fluorescence contributed by the B-C6 band divided by the cumulative fluorescence of all bands in each TLC lane; ****p < 0.0001. **(D)** Apical-reservoir fluorescence of B-C12, or B-C16 treated AE Transwell cultures treated with vehicle (DMSO) or etomoxir; * p < 0.05, ****p < 0.0001. **(E, F)** Left, cellular-reservoir fluorescence of B-C12, or B-C16 treated AE Transwell cultures treated with vehicle (DMSO) or etomoxir; ** p < 0.01, ****p < 0.0001. **(G)** Schematic showing proposed mechanism of reduced B-FA export by etomoxir. B-Cx denotes fluorescence coming from all B-lipids in basal, apical or cellular reservoirs. A previous study^62^ using TLC to identify B-lipid species with the same solvents used in this study reported naturally fluorescent bands that do not correspond to B-lipid standards used. Approximate locations of NFBs are labelled as NFB. *Location of B-C4, B-C6 and B-C8 were inferred from the experiment in supplemental figure 4. B-FAs were applied at a concentration of 20 uM. Etomoxir was applied at a concentration of 100uM. Experiments were performed in triplicate (n = 3 AE transwell cultures).

Strong barrier function through the duration of B-FA and etomoxir treatment was confirmed by lack of Dextran-647 in the basal reservoir (Fig. S5). When B-C5 was the apical input FA, basal fluorescence did not significantly change when FAO was inhibited (Fig. 5A). By contrast, when B-C12 and B-C16 were the apical input FAs, FAO inhibition significantly reduced basal-reservoir fluorescence (Fig. 5A). TLC demonstrated that the most abundant lipid species in the basal reservoir of B-C5 treated cultures was B-C5 (Fig. S6), suggesting that this FA passively diffused through AE monolayers, as FAs become increasingly water soluble as chain-length decreases^65^. For control cultures (vehicle) treated with B-C12 and B-C16, the most predominant lipid species corresponded to B-C6 (Fig. 5B,C), indicating catabolism of B-C12 and B-C16.

Following treatment of B-C12 and B-C16 cultures with the FAO inhibitor, etomoxir, B-C6 significantly decreased (Fig. 5B,C). In B-C12 treated cultures, etomoxir caused the apical input FA (B-C12) to be the predominant lipid species in the basal reservoir (Fig. 5B). In cultures exposed to apical B-C16, etomoxir treatment did not result in B-C16 being the predominant lipid species in the basal reservoir (Fig. 5C). These findings suggest that long-chain FAs (> B-C12) are less amenable to passive diffusion than short-(B-C5) and medium-(B-C12) chain FAs. Together these data indicate that FAs of shorter-chain length (B-C5, B-C6) are more amenable to basal export by AEs than FAs of longer-chain length (B-C12, B-C16), which appear to require catabolism via FAO to generate a shorter FA derivative (B-C6) that can then be readily exported or diffuse to the basal reservoir.

While total basal-reservoir fluorescence and TLC show that etomoxir reduces FA export in B-C12 and B-C16 cultures, these results cannot rule out the possibility that etomoxir reduces basal-reservoir fluorescence by inhibiting apical FA import thus resulting in less intracellular FA available to undergo FAO. To test whether etomoxir reduced apical FA import, fluorescence from the apical reservoir of B-C12 and B-C16 treated cultures was quantified following 6-hours of etomoxir treatment (Fig. 5D). If etomoxir reduced FA import, there would be more total B-FA fluorescence in the apical reservoir compared to control, however, the data demonstrated the opposite, that etomoxir significantly reduced apical reservoir fluorescence in cultures exposed to B-C12 and B-C16 (Fig. 5D). These results support the conclusion that etomoxir does not reduce apical FA uptake.

The finding that etomoxir significantly reduced both apical and basal reservoir fluorescence was somewhat surprising and raised the hypothesis that impaired FAO resulted in accumulation of B-FAs in the cellular reservoir. To test this, AE monolayers were treated with B-C12 or B-C16 along with vehicle or etomoxir. Following 6 hours of apical exposure, AE monolayers were imaged to quantify cellular fluorescence (Fig. 5E,F). Etomoxir significantly increased cellular reservoir fluorescence in B-C12/16 treated cultures, further supporting the conclusion that impaired FAO reduced FA export (Fig. 5E). Together, these data support the conclusion that medium- and long-chain FAs are catabolized by FAO to smaller-chain FAs that can be exported as free FAs (Fig. 5G).

### Metformin and C75 potentiate FAO in AE monolayers and increases basolateral export of oxidized FAs

To further support a role for FAO in regulating basal export of B-FAs, it was hypothesized that treating AE monolayers with drugs that augment FAO would result in increased FA export. To test this hypothesis, AE monolayers were independently treated with metformin and C75, drugs that demonstrate FAO augmentation in other cell types^66,67^. Metformin, commonly known as an anti-diabetic drug, potentiates FAO by preventing formation of the CPT1 inhibitor malonyl-CoA^68^ and C75, a weight-loss-inducing drug, potentiates FAO by increasing CPT1 activity^69^ (Fig. 6E). Following metformin or C75 exposure, B-C12 or B-C16 were applied to the apical reservoir to mimic post-prandial FA exposure. Media from the basal reservoir was collected at 2-hour intervals for 6 hours and total fluorescence was quantified to measure FA export. Metformin and C75 significantly increased fluorescence in the basal reservoir of B-C12 and B-C16 treated monolayers (Fig. 6A). TLC was performed to determine the lipid species exported to the basal reservoir of B-C12 and B-C16 cultures (Fig. 6B,C). A lipid species corresponding to B-C6 was the most abundant B-FA species in the basal reservoir of B-C12 and B-C16 treated cultures (Fig. 6B,C) indicating that B-C6 is the primary B-FA metabolite exported.

**Figure 6.**
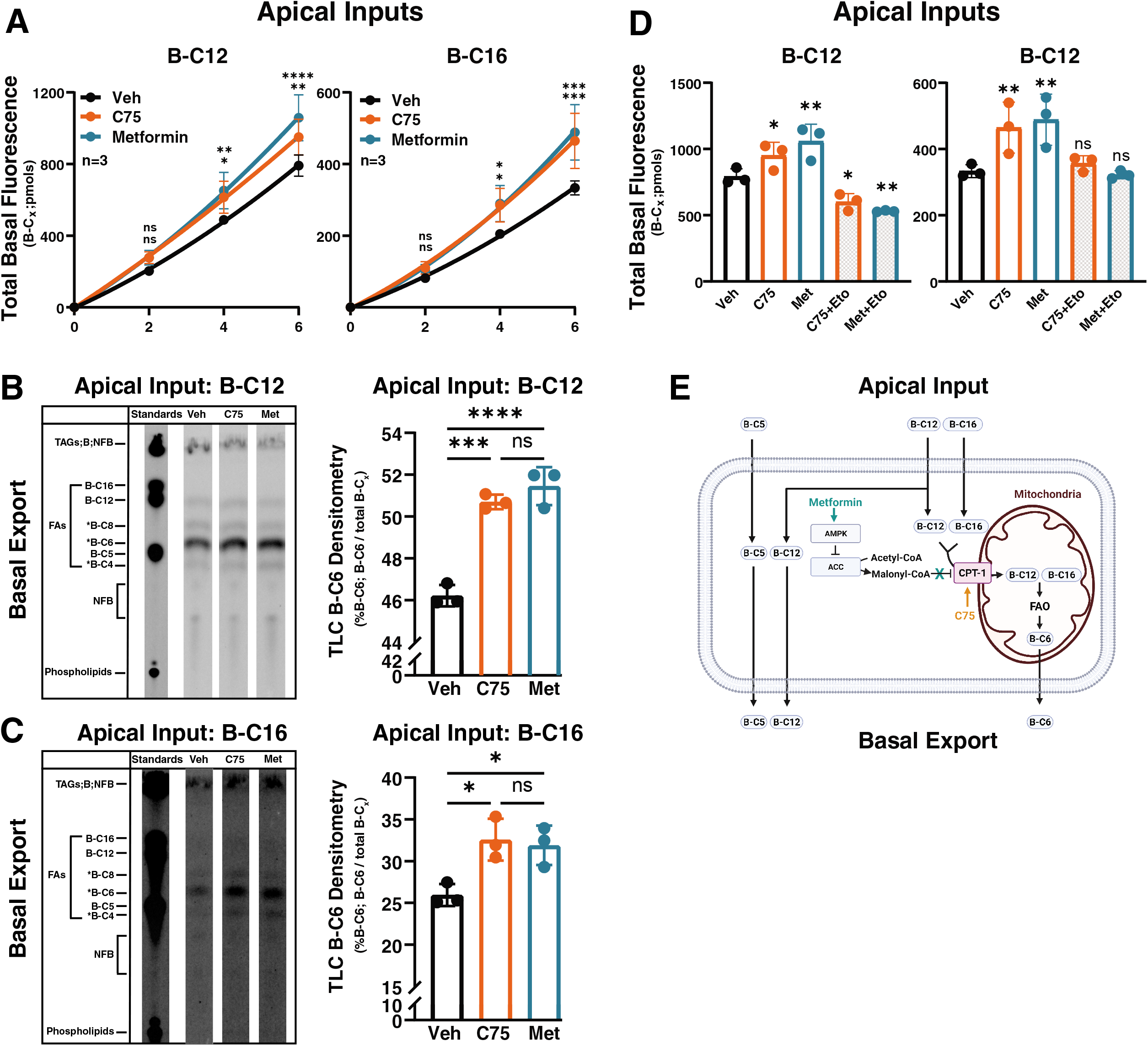
Enhancing FAO augments FA export. **(A)** Quantification of basal-reservoir fluorescence from B-C12 or B-C16 treated AE cultures treated with vehicle (DMSO), C75, or metformin (Met); * p < 0.05, ** p < 0.01, *** p < 0.005, **** p < 0.0001. **(B, C)** Left, TLC of basal-reservoir media from B-C12 or B-C16 treated AE cultures treated with vehicle, C75 or metformin.; * p < 0.05, *** p < 0.001, **** p < 0.0001. **(D)** Basal-reservoir fluorescence of B-C12 treated cultures treated with either vehicle, C75, metformin, C75 and etomoxir, or metformin and etomoxir. Fluorescence was measured 6-hours post B-FA application; * p < 0.05, ** p < 0.01. **(E)** Proposed mechanism of enhanced FA export by C75 and metformin. Concentrations of C75, metformin and etomoxir were 40uM, 3mM, and 100uM respectively. A previous study^62^ using TLC to identify B-lipid species with the same solvents used in this study reported naturally fluorescent bands that do not correspond to B-lipid standards used and are labelled here as NFB. *Location of B-C4, B-C6 and B-C8 were inferred from the experiment in supplemental figure 4. Experiments were performed in triplicate (n = 3 AE transwell cultures).

To probe whether the B-FA export effects were dependent on FAO, AE monolayers exposed to B-C12 or B-C16 were co-treated with each of the FAO potentiators, C75 or metformin, and the FAO inhibitor, etomoxir. After 6hrs of B-FA exposure and drug treatment, basal reservoir fluorescence was quantified. Etomoxir blocked the ability of C75 and metformin to increase basal reservoir fluorescence in cultures exposed to B-C12 and B-C16 (Fig. 6D), supporting the conclusion that metformin and C75 function in an FAO-dependent manner to increase basal export of oxidized long- and medium-chain FA metabolites.

## Discussion

Once born from an ISC, progenitor cells transition through a number of lineage states during their 7-day lifespan *in vivo*^70^. Mouse studies suggest the absorptive lineage transitions through functional maturation states classified as early-, intermediate-, and mature-phases as they migrate up the villus axis^50^. These functional maturation states are associated with different biological roles related to antimicrobial functions early-on and progress into nutrient handling function toward their late and terminal maturation stage^50^. Here we demonstrate by single-cell transcriptomics that human jejunal AEs *in vivo* generally share a similar maturation defined by early-, intermediate- and late-maturation phases. Importantly, these states correlate with discrete lipid-handling gene profiles as AEs mature. While our study focused on lipid-handling genes and mechanisms, the transcriptomic datasets will be useful to evaluate aspects of nutrient handling (*i.e.,* carbohydrate, protein, and vitamin metabolism) in primary human AEs.

The absorptive lineage is the default pathway taken by progenitor cells unless the master regulator of secretory lineage fate, Atoh1, is expressed^55^. Chromatin states across the genome are associated with hardwiring of some lineage-specification programs^71,72^, and dynamic chromatin states have been described as cells move through ISC to secretory and absorptive lineage states^71,72^. Data presented here demonstrate that removal of ISC maintenance factors from ISC monolayers promote a stereotypical AE lineage program in an epithelial autonomous manner suggesting human AEs are hardwired to undergo progressive maturation over time. Since our *in vitro* AE monolayer system generally mimics *in vivo* AE maturation and lifespan, it is highly suited to define the intrinsic nature of chromatin dynamics through human AE maturation and is amenable to testing how extrinsic influences such as dietary factors or the microbiome might influence chromatin states and associated gene expression.

The majority of dietary lipids consist of LCFAs. LCFAs undergo intracellular esterification to TAGs, are packaged into chylomicrons, exported through the basal membrane, and distributed through the lymphatic system^1,2^. Previous studies have demonstrated that SCFAs and MCFAs can bypass the chylomicron-lymphatic pathway and pass unesterified into the portal vein^73^. Because LCFAs can be oxidized to SCFAs and MCFAs via FAO, we hypothesized that FAs derived from FAO of LCFAs could be exported through the basal membrane as free FAs. Using our culture system, we demonstrate for the first time that smaller-chain FAs generated from FAO of LCFAs can be exported unesterified across the basal membrane of AEs. We demonstrate a dependency on FAO in regulating basal export of LCFA derived FAs as their basal export was decreased by inhibiting FAO with etomoxir and increased by enhancing FAO with C75 and metformin. This novel pathway of LCFA derived FA export might explain why patients with abetalipoproteinemia exhibit distribution of the majority of dietary FAs yet are unable to secrete chylomicrons^74^.

Basal export of free FAs regulated by FAO could be involved in other physiological responses to dietary lipids, disease etiologies, and pharmaceuticals that target FAO pathways. Our findings demonstrate that metformin, a satiety-inducing/glucose-lowering drug, enhances basal FA export, raising the possibility that FAO and augmented basal FA export underlies some of metformin’s efficacies. In this regard, stimulation of free fatty acid receptors (FFARs), which are restricted to the basal surface^75^ of enteroendocrine cells (EECs)^69^ immediately adjacent to AEs, stimulate secretion of glucose-lowering GLP-1 and satiety-inducing PYY^76^ gut hormones. Reports demonstrate that metformin elicits intestinal GLP-1 secretion by EECs and that this effect significantly contributes to systemic glucose-lowering^77^.

Importantly, exposure of metformin to immortalized EEC lines fails to stimulate GLP-1 secretion suggesting metformin does not act directly on EECs to stimulate GLP-1 secretion but rather through a more complex mechanism^78^. In light of our results demonstrating metformin increases FA export, metformin may act to increase GLP-1 secretion via an AE-FAO-EEC axis. In this scenario apically localized MCFAs and LCFAs are absorbed by AEs, catabolized to shorter-chain FA species that can passively diffuse through the basolateral membrane where they interact with nearby FFARs on EECs to stimulate GLP-1 release. Further development of our AE culture system to support co-culture of EEC and AEs with loss- and gain-of-function for key FAO genes will be required to test this hypothesis.

Aside from basic science applications, our high-throughput 96-well AE culture platform is highly suitable for drug screening and validation. Compared to standard 12-well Transwell plates, scaling the platform to 96-wells increases the plate form factor by 8 while simultaneously requiring approximately 8-times less cells. This substantially increases the number of biological and technical replicates that can be performed per plate and reduces plate-to-plate variability when performing experiments to evaluate therapeutic indices for a drug. Real-time quantification of FA mobilization through the epithelial barrier by plate reader allows for kinetic studies while the planar format of these cultures facilitates simultaneous high-content microscopic readouts. While the AE monolayer surface and reservoir sample volumes are small, we show there remains sufficient material for RNA-sequencing and highly sensitive detection of B-conjugated FAs and metabolites by plate reader and TLC. The scalability, physiological relevance, and sensitivity of our platform to detect changes in FA-handling could facilitate the discovery of treatments for metabolic disorders impacted by intestinal FA-handling such as obesity, dyslipidemia, and diabetes.

## Materials and Methods

### Donor Selection

Human donor intestines were accepted and received from HonorBridge (formerly Carolina Donor Services, Durham, NC) based on the following donor acceptance criteria: age ≤ 65 years, brain-dead only (as opposed to donation after cardiac death), negative for HIV, Hepatitis, RPR (syphilis), tuberculosis, or COVID-19. Tissue from a 29-year-old Caucasian male was used for single cell dissociation and scRNAseq. Tissue from a 51-year-old African American male was used for tissue culture, *in situ* hybridization, scRNAseq of collagen grown ISCs and AE monolayers and bulk RNAseq. Human donors had no history of bowel surgery, severe abdominal injury, cancer, or chemotherapy. Increased risk donors (*i.e.,* history of incarceration or intravenous drug use) were accepted, provided negative infectious disease results. Additionally, donor cases where the pancreas was placed for transplant were excluded given that pancreatic transplants require removal of proximal small intestinal tissue.

### Organ resection and single cell dissociation

Whole human intestines were transported to UNC Chapel Hill in ice-cold University of Wisconsin Solution, with tissue dissection beginning within eight hours of cross-clamping. First, fat, and connective tissue were trimmed from the donated organs and intestines were subdivided into six regions following measurement. For the small intestine, the proximal 20 cm was deemed Duodenum. Jejunum and ileum were determined through an even split of the remaining small intestine. Two 3×3 cm^2^ resections were isolated from the center of jejunum and ileum for dissociation.

Resections were incubated in 10 mM NAC at room temperature for 30 min to remove mucus, then tissue was moved to ice-cold Isolation Buffer which consisted of 5.6 mM Na2HPO4, 8.0 mM KH2PO4, 96.2 mM NaCl, 1.6 mM KCl, 43.4 mM Sucrose, 54.9 mM d-sorbitol, and 100 uM Y27632, then washed several times by gently inverting the tubes. Tissues were then incubated in Isolation Buffer with 2 mM EDTA and 0.5 mM DTT, then shaken vigorously to remove crypts. Shakes were repeated several times, checking for crypts and/or villi each time. High-yield small intestinal shakes were pooled to approximate 1:1 villus to crypt tissue by cell mass. Crypts and villi were dissociated to single cells using 4 mg/ml Protease VIII in DPBS + Y27632 on ice for ~45min with trituration via a P1000 micropipette every 10 min. Dissociation was checked on a light microscope then clumps were removed using filtration.

### Cell sorting, library prep, and sequencing

Single cells were washed with DPBS + Y27632 then resuspended in Advanced DMEM/F12 + 1% Bovine Serum Albumin + Y-27632. AnnexinV-APC (1:100) was added for live/dead staining and one TotalSeq Anti-Human Hashtag Antibody per region to allow for tracking all six regions with a single library preparation. Cells were washed with Advanced DMEM/F12 + 1% BSA +Y27632 then resuspended in the same solution for sorting on a Sony Cell Sorter SH800Z. Cells were gated using forward and backward scatter and AnnexinV to enrich for live single epithelial cells. 25k cells were collected from each separate region, then all regions were combined before sequencing. Library prep was performed with the Chromium Next GEM Single Cell 3’ GEM, Library & Gel Bead Kit v3.1. Sequencing was performed on an Illumina NextSeq 500.

### Single cell RNA sequencing data processing

After sequencing, reads were aligned to reference transcriptome GRCh38 with the 10X Cell Ranger pipeline. Mapped reads were filtered and counted by barcode and UMI and then transformed into an AnnData object using the Python implementation of scanpy (v1.7.2). Annotations for cell cycle phase were added following previously published methods^79^. The number of genes, number of UMIs, and percent mitochondrial expression for all cells in each sample were visualized and used to identify thresholds for high-quality cells to include in further analysis(<u>Fig. S1F,G and S7A,B</u>).^80^. Quality control parameters for both datasets are:

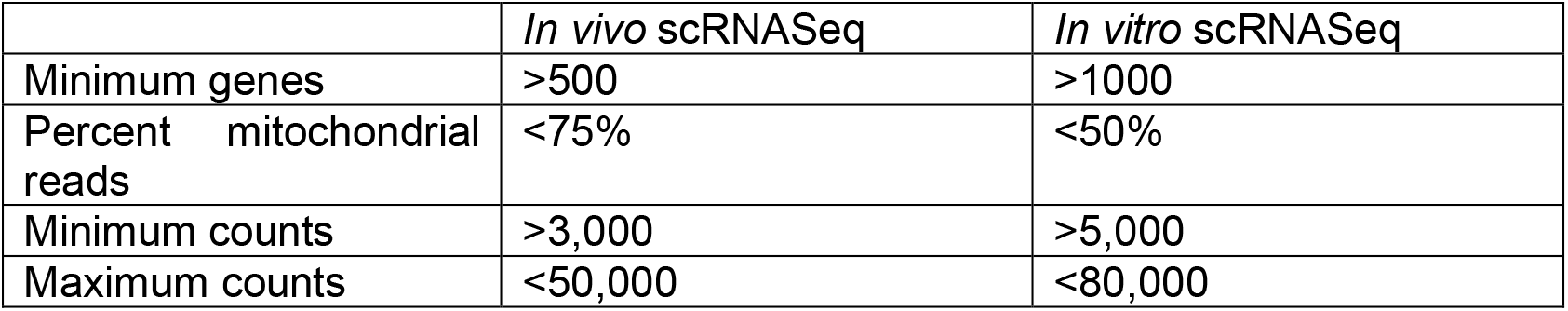

Following filtering, read counts were log-transformed and normalized to the median read depth of the dataset. For both single cell sequencing experiments no batch correction was performed as each dataset was analyzed separately. Variability due to read count, percentage of mitochondrial reads and cell cycle phase were regressed out by simple linear regression. Highly variably genes were identified using Seurat v2. 2,585 and 4,186 highly variable genes were used for principal component analysis for the *in vivo* and *in vitro* scRNAseq datasets, respectively. Counts for each gene were scaled to have a mean of zero and unit variance. A kNN-graph was constructed with 10 neighbors was used to calculate Leiden clusters for both *in vivo* and *in vitro* scRNAseq datasets (*In vivo* clustering parameters: Leiden resolution = 0.5, num_neighbors = 10, num_pcs = 40; *in vitro* clustering parameters: Leiden resolution = 0.1, num_neighbors = 10, num_pcs = 15) and EPCAM-negative cells were removed from the *in vivo* dataset^81^. PAGA was used to initialize UMAP embeddings of Leiden clusters^82^.

Regional hashtag deconvolution followed published methods for both scRNAseq datasets. Briefly, raw hashtag read counts were normalized using centered log ratio transformation followed by k-medoid clustering (k=6 medoids). Hashtag noise distributions were determined by removing the cluster with highest expression of a specific hashtag, then a negative binomial distribution was fit to the data of the remaining cells. Cells were considered positive for a hashtag if counts for the specific hashtag were above the distribution’s 99^th^ percentile (p<0.01) threshold. Cells positive for multiple hashtags were excluded as likely doublets. Cells called for colon and duodenum were removed from the dataset. For *in vitro* data, only hashtags 1 and 3 were kept for analysis.

### RNA Velocity

Velocyto 0.17.16 was used to generate the initial loom file and scvelo 0.2.4 was used for all integration of spliced/unspliced loom file integration with the processed anndata object and all subsequent trajectory analysis^51,52^. Briefly, to ensure connectedness of dataset, clusters that corresponded to stem cells and the different stages of enterocyte differentiation were used for trajectory analysis. All other clusters were removed from the dataset and the remaining clusters were reprocessed and reclustered. The resulting dataset was integrated with the spliced and unspliced read counts and 4100 highly variable genes were kept for fitting to the RNA velocity dynamical model. For moment calculation, num_pcs = 40 and num_neighbors = 50. Function arguments for calculating RNA velocity vectors and latent time were default values except for the specification of the dynamical model in scvelo.tl.velocity.

### Differential Gene expression for scRNASeq

Differential expression analysis was performed using the de.test.wald function in the Python implementation of diffxpy (version 0.7.4)^83^. From the output data, significant DEGS (q <0.05) were attributed to the cluster with the highest mean value for each gene.

### Linear correlation Analysis

Read count data from both datasets were normalized to the same value then log-transformed. Mean expression values for each lipid-handling gene were calculated per-scRNAseq cluster and per group of *in vitro* differentiation bulk RNAseq. Pairwise comparisons of mean gene expression values for each *in vivo* differentiation state and each *in vitro* differentiation state were made (Fig. S4B). A line was drawn representing a perfect correlation of *in vitro* bulk RNAseq expression to *in vivo* scRNAseq expression (Fig, S3B). Residuals for each lipid-handling gene were calculated based on the deviation from this line. The residual sum of squares thus represents a quantitative measure of the overall similarity of expression of lipid-handling genes between each *in vivo* differentiation state and each *in vitro* differentiation state, with a lower value indicating a better fit to the line describing a perfect correlation.

### Transwell Preparation, ISC Seeding, Expansion and Differentiation

The apical surface of 12-well and 96-well permeable Transwell inserts was overlayed with ice cold 1% Growth Factor Reduced Matrigel (Matrigel) diluted in ice cold DPBS. Transwell plates were left incubating at 37C in 5% CO2 overnight. Inserts were rinsed by replacing the 1% Matrigel with DPBS the following day. Jejunal ISCs grown on 2D collagen scaffolds were suspended in EM containing 10 mM Y27632 using an established protocol^38^. Suspended ISCs were seeded on the apical surface of Transwell inserts at densities of approximately 300K-400K cells/cm^2^ (Table S1). EM was added to the basal reservoir at the time of seeding. Apical and basal media was replaced the day after seeding with fresh EM (Table S2). EM in apical and basal reservoirs was replaced with DM to initiate differentiation (Table S2).

### Single cell dissociation of Transwell epithelial monolayers

Media was removed from ISCs expanded on collagen and from AEs on 12-well transwell inserts and washed with 1x DPBS. 1.5mls of 3mM EDTA in PBS was applied to ISCs expanded on collagen and on apical and basal reservoirs of 12-well transwell cultures until most cells were lifted off the collagen or Transwell surface as determined by visual inspection every 2 minutes under an inverted microscope. EDTA was aspirated using a P1000 micropipette and redistributed over the Transwell surfaced to facilitate detachment of cells from the Transwell at 2-minute intervals until the majority of cells had detached. After the cells detached, 1.5mls of DPBS was applied to each well and rinsed by aspirating and re-applying the EDTA/DPBS. Cells were then aspirated and pelleted by centrifugation at 500g for 5 minutes. The supernatant was removed and replaced with 1ml of 4mg/mL cold protease in DPBS. Cells in cold protease were incubated on ice and pipetted every 2 minutes before visual examination of dissociation under an inverted light microscope. This was repeated until all cells were singlets. Following single cell dissociation, the cold protease was quenched with Advanced DMEM/F12 + 1% FBS. Cells were pelleted as described above and resuspended in Advanced DMEM/F12 +1% FBS.

### Bulk RNA sequencing preparation, processing, and analysis

To investigate the dynamic changes in gene expression as ISCs differentiate into AEs *in vitro*, RNA-seq was performed on human intestinal epithelial monolayers immediately before seeding onto Transwells (D0) and at Days 2, 5, 7, 10, and 11 (D2, D5, D7, D10, D11) of differentiation on Transwells. N=3 samples were collected from each time point and RNA was extracted using RNAqueous-Micro Total RNA Isolation Kit according to manufacturer’s protocols and stored at −80°C. RNA quality was assessed prior to library preparation by using the Agilent 2100 Bioanalyzer to determine the RNA integrity number (RIN)^84^. After confirmation that each sample had a RIN of at least 8, integrated fluidic circuits (IFCs) for gene expression and genotyping analysis were prepared using the Advanta™ RNA-Seq NGS Library Prep Kit for the Fluidigm Juno™ and sequenced with the Fluidigm Biomark™ HD system. Gene level expression was obtained through pseudo alignment of reads to human genome GRCh38 using Kallisto^85^. Expression values for plotting were obtained by TMM normalization across all samples using EdgeR package^86^. Sequencing data from each time point were combined and then were normalized to the dataset median and log-transformed. Principal component analysis was done with scikitlearn (v0.24.0).

### Transepithelial Electrical Resistance

Barrier integrity of 96-well Transwell cultures was monitored by quantifying TEER using EVOM or EVOM3 TEER meters in conjunction with STX100C96 electrodes.

### Basal Fluorescence Quantification

100uM Etomoxir, 40uM C75, 3mM metformin or 1x vehicle (DMSO) were suspended in 100uls of DM and applied to the apical surface of AE Transwell cultures for 1 hour. Following incubation, apical media was replaced with 100ul of DM containing drugs or vehicle, 1uM Dextran, Alexa Fluor 647 (Dextran-647) and 20uM of either BODIPY-FL C5, −C12 or −C16. Basal media (50ul) was collected at 2, 4 and 6 hours and replaced with an equivalent volume of DM. Quantification of basal fluorescence was performed using a CLARIOstar Plus Microplate Reader. Fluorescence arbitrary units (AU) were converted to pmols using a standard curve.

### Thin Layer Chromatography

For the assessment of FA-handling via TLC (Fig. 4), BODIPY-FL C12 was applied to the apical surface of AE Transwell cultures. Apical and basal media was collected after 6 hours. Cells were released from Transwell inserts via application of 100ul of 1x TrypLE Express containing 10mM Y27632. Cells were lysed by undergoing 3 freeze-thaw cycles. Lipid extracts were generated from apical and basal media along with cell lysates using the Bligh and Dyer method ^87^. Lipid extracts were resuspended in 100% ethanol and spotted on silica gel TLC plates. Lipid species were separated by placing spotted silica gel plates in a glass chamber containing either a polar (chloroform/ethanol/triethylamine/water, 30:34:30:8 mL) or protonating (petroleum ether/ethyl ether/acetic acid, 30:34:30:8 mL) solvent. The following BODIPY analogs were used to identify lipid species BODIPY-FL C5 (B-C5), −C12 (B-C12), −C16 (B-C16), BODIPY 493/503 (B) and β-BODIPY-FL C12-HPC (phospholipids). BODIPY-FL C4 (B-C4), −C6 (B-C6) and −C8 (B-C8) are not commercially available therefore their location on TLC plates was inferred by generating a standard curve of the distances travelled by BODIPY-FL C5, −C12 and −C16 in polar solvent. 18:1-18:1-C11 TopFluor TG is a TAG conjugated to a fluorophore with similar properties (chemical structure and excitation/emission) as BODIPY and was used to infer the location of TAGs on TLC plates due to BODIPY-TAG analogs not being commercially available at the time of this publication. Only basal media samples were collected and assessed via TLC from the experiments in figure 5 and figure 6. Following incubation in solvent, spotted silica gel plates were dried and scanned on an iBright FL 1000 Imager to detect fluorescent lipid species. Excitation and emission channels were set to 455-485 and 508-557 respectively. A previous study^62^ using TLC to identify B-lipid species with the same solvents used in this study reported naturally fluorescent bands that do not correspond to B-lipids and are labelled here as NFB.

### Microscopy and Image Analysis

AE monolayers were treated with vehicle or 20uM etomoxir and exposed apically to B-C12 or B-C16 for 6 hours. After 6 hours of apical B-C12 or B-C16 exposure, AE monolayers were rinsed with fresh DM and fixed in 40% glyoxal solution for 20 minutes. Following fixation, AE monolayers were rinsed with 1x DPBS. Next, the transwell membrane containing the fixed AE monolayers were removed from the transwell inserts and placed on glass slides. AE monolayers were then overlayed with mounting media and covered with a glass coverslip. Fluorescent images of AE monolayers were taken at 40x magnification on a Keyence BZ-X810 microscope.

## Code and Data Availability

Sequencing datasets will be available on the NCBI Gene Expression Omnibus under accession number GSE186583. Python scripts demonstrating the main parts of our analysis will be available on GitHub.

## Supplemental Figure/Table Legends

**Supplemental Table 1**

Reagents, materials, and instruments

**Supplemental Table 2**

Media conditions

**Supplemental Table 3**

DEGs of jejunal ISC, eAE, iAE, and mAE lineages as determined by DEG analysis comparing these populations.

**Supplemental Figure 1.**
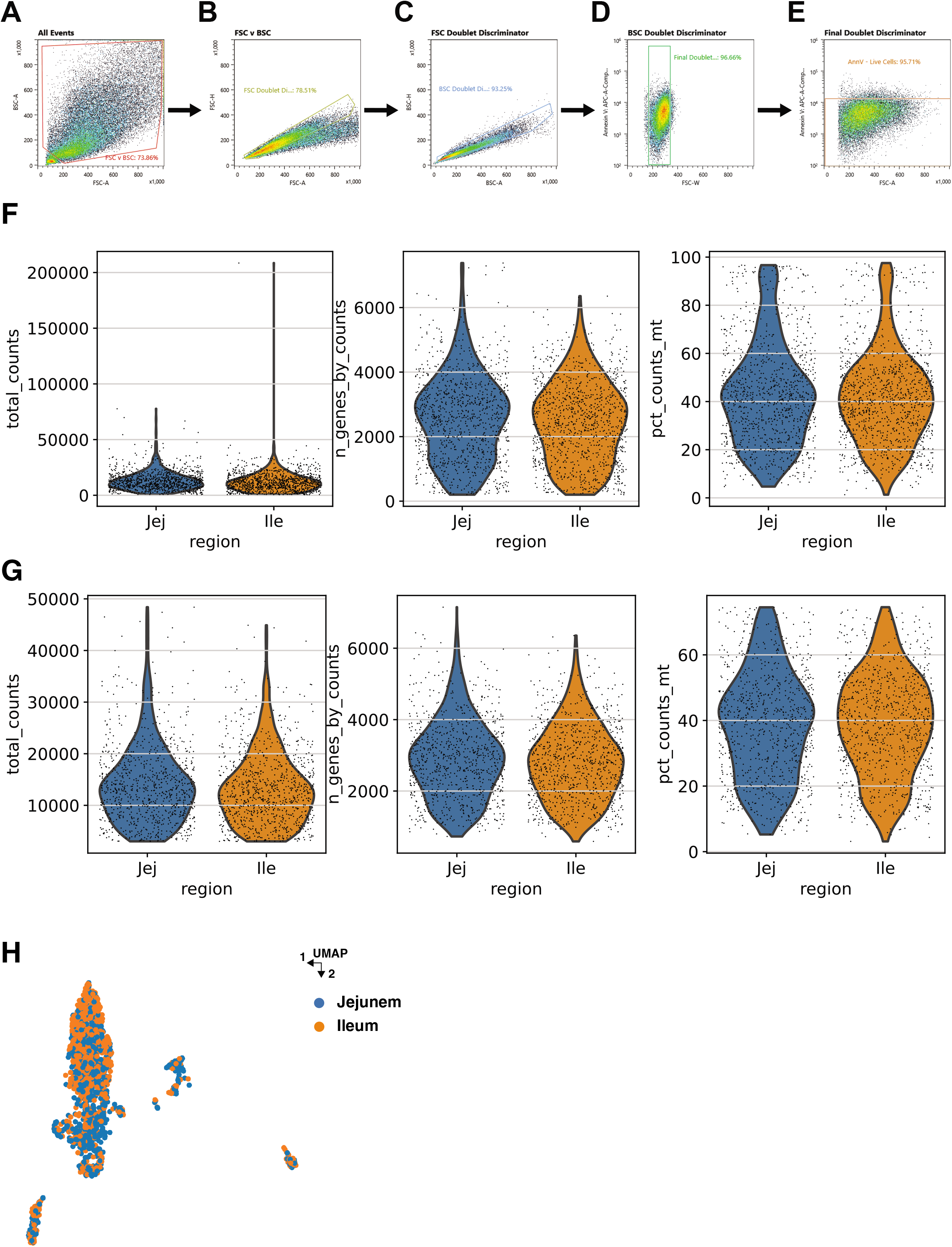
**(A-E)** FACS density plots showing FACS gating strategy for sorting live cells from dissociated human small intestine. Titles above plots indicate the gate which the cells came from (*i.e.,* the previous density plot). **(A)** shows initial doublet discriminator, with exclusion of likely red-blood cells and immune cells. **(B)** Forward-scatter based doublet discriminator. **(C)** Back-scatter only doublet discriminator. **(D)** Forward-scatter width-based doublet discriminator. **(E)** Final gate used to distinguish live cells based on negative gating for AnnexinV-APC. **(F,G)** Violin plots showing distributions of QC parameters including: number of total reads per cell, number of genes counted in each cell, and the percent mitochondrial reads per cell. **(F)** Pre-filtering distribution of QC parameters. **(G)** Post-filtering distribution of QC parameters. **(H)** Region of cells overlayed on UMAP.

**Supplemental Figure 2.**
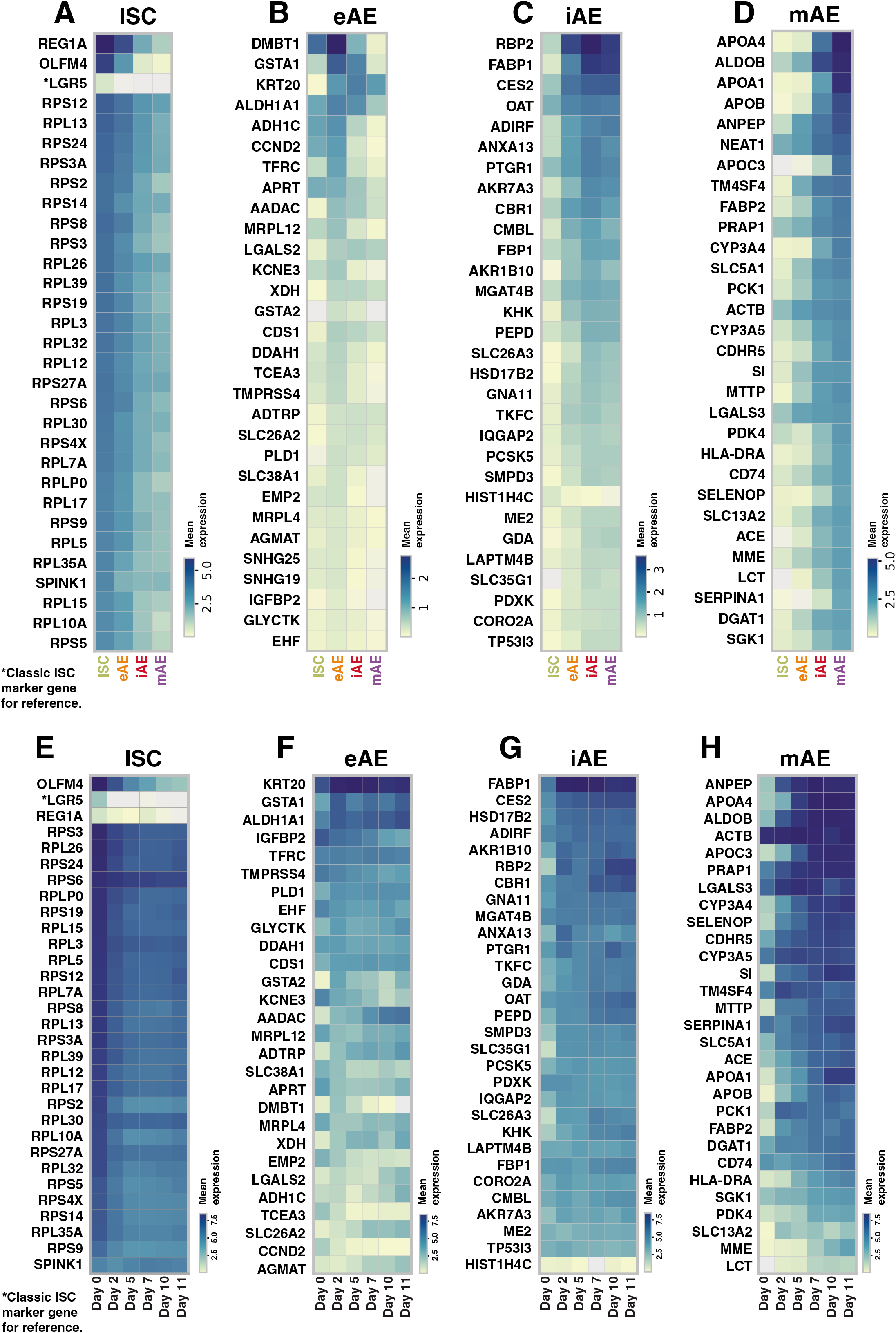
Expression of top 30 DEGs in ISCs **(A)**, eAEs **(B)**, iAEs **(C)**, and mAEs **(D)**. * LGR5 was not among the top 30 DEGs in ISCs but it was included for reference. *In vitro* expression of top 30 *in vivo* ISC **(E)**, eAE **(F)**, iAE **(G)** and mAE **(H)** DEGs. *SNHG19, SNHG25* and *NEAT1* were not detected in our *in vitro* data set and were excluded.

**Supplemental Figure 3.**
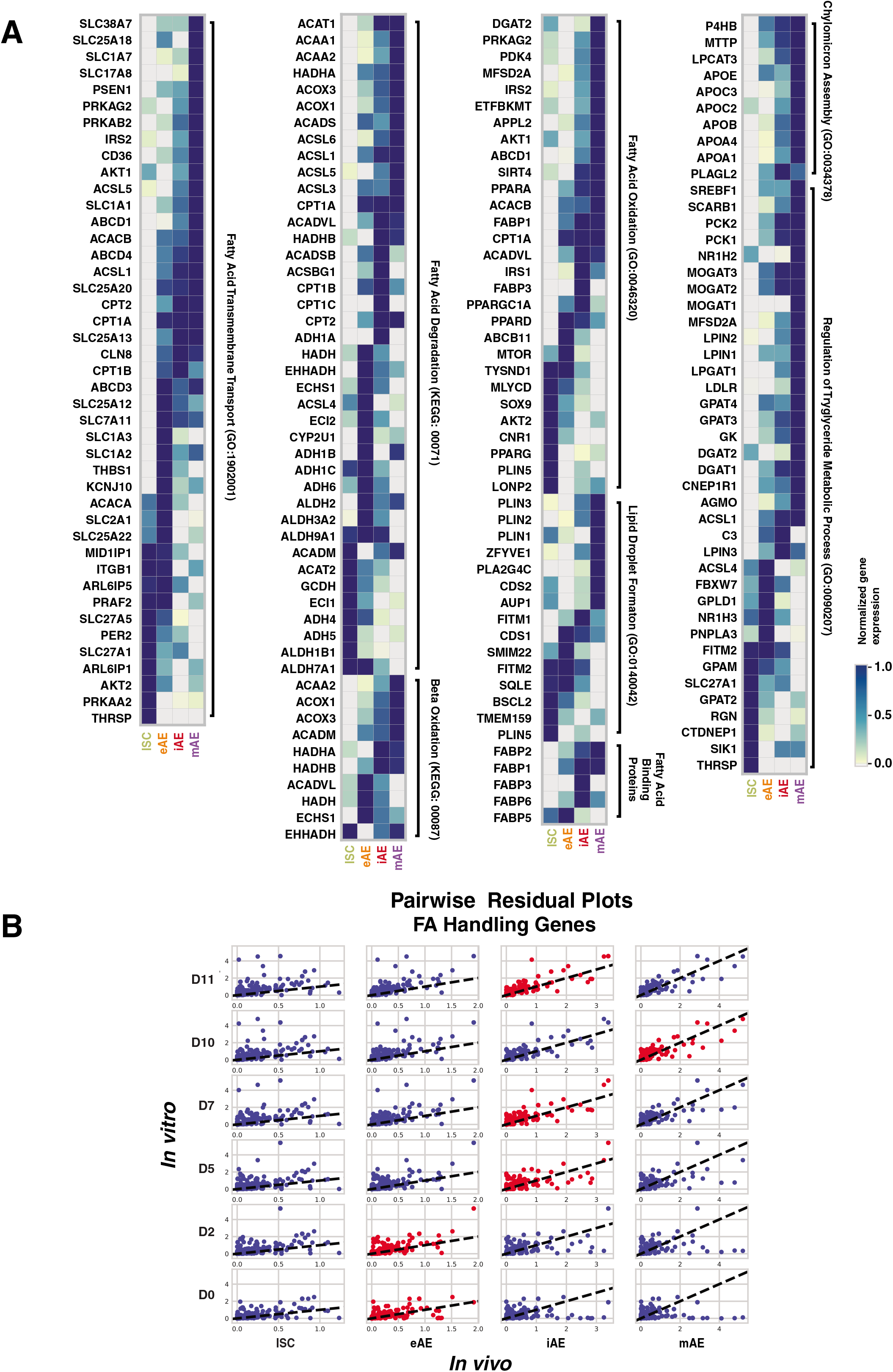
**(A)** Expression of lipid-handling genes in *in vivo* ISC and human AE populations. Genes were selected from Go terms relating to lipid metabolic processes. Fatty acid binding protein genes were selected manually and not derived from Go terms (*bottom* left). The following genes were not detected in our dataset and were excluded; *DGAT2L7P, MIR29B1, MIR30C1, MIR548P, APOA2, PLIN4 ABCD2, GRM1, NTSR1, SLC17A6, SLC17A7, SLC1A6, ABCD2, TWIST1*. **(B)** Matrix showing linear regression plots for each gene described in figure 3C. Each dot represents a gene from the above matrix plots. The dotted line shows the line that was used to calculate residuals and is drawn with slope = 1 (*i.e., in vitro* differentiation bulk RNA seq (n=3 samples per time point) mean expression perfectly matches *in vivo* scRNAseq mean expression for each cluster). Red dots indicate lowest residual sum of squares for each row (*i.e.,* lowest residual for each time point sampled in the *in vitro* differentiation experiment).

**Supplemental Figure 4.**
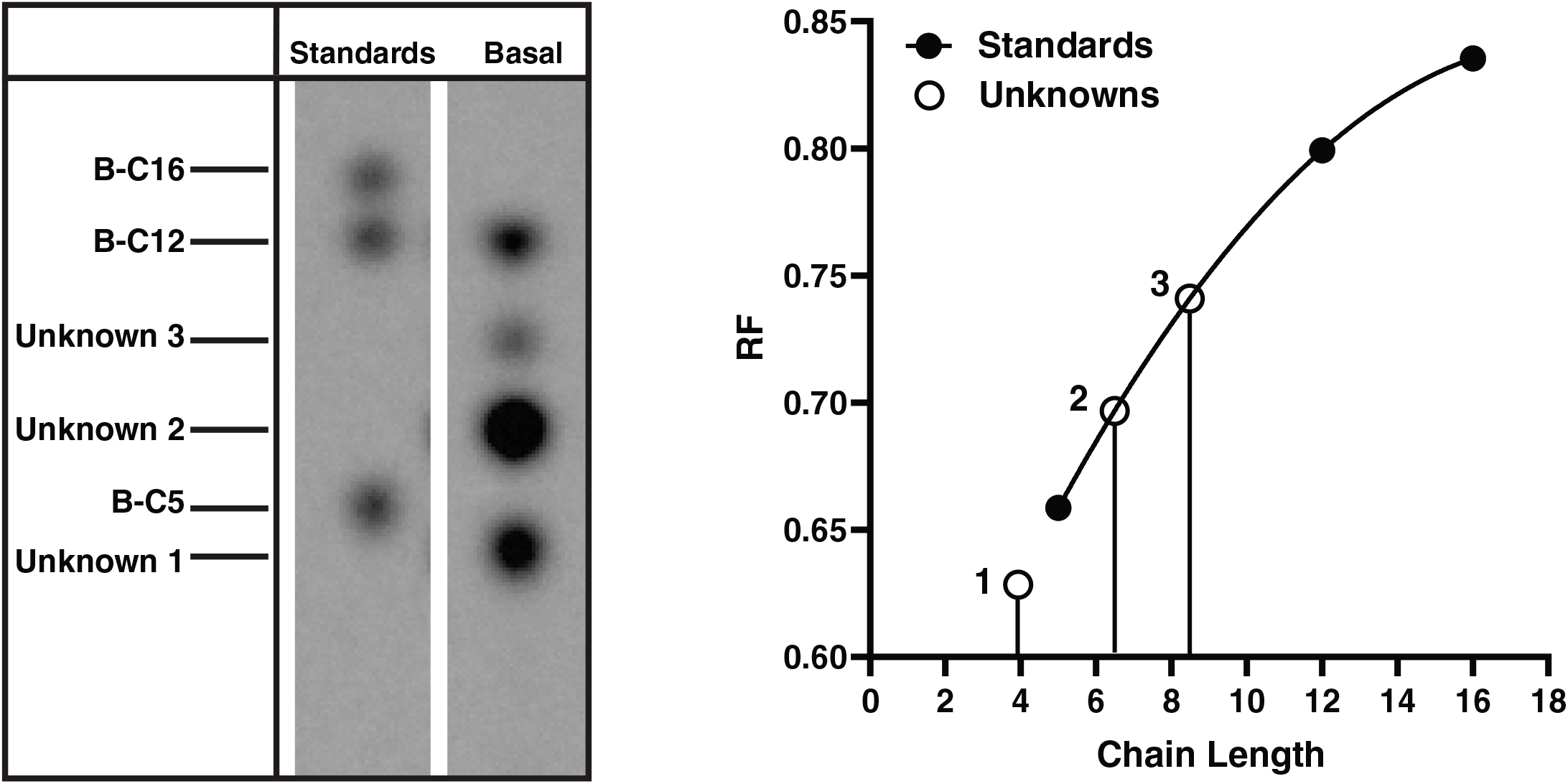
TLC (polar) of basal media from AE cultures after 6 hours of incubation with B-C12 (Left). Retention factor (Rf) of B-C5, −C12 and C16 standards were used to generate a standard curve to infer the chain length of FAs whose Rf values do not correspond to B-FA standards used (right). The three unknown FAs are predicted to correspond approximately to B-C4,-C6 and −C8.

**Supplemental Figure 5.**
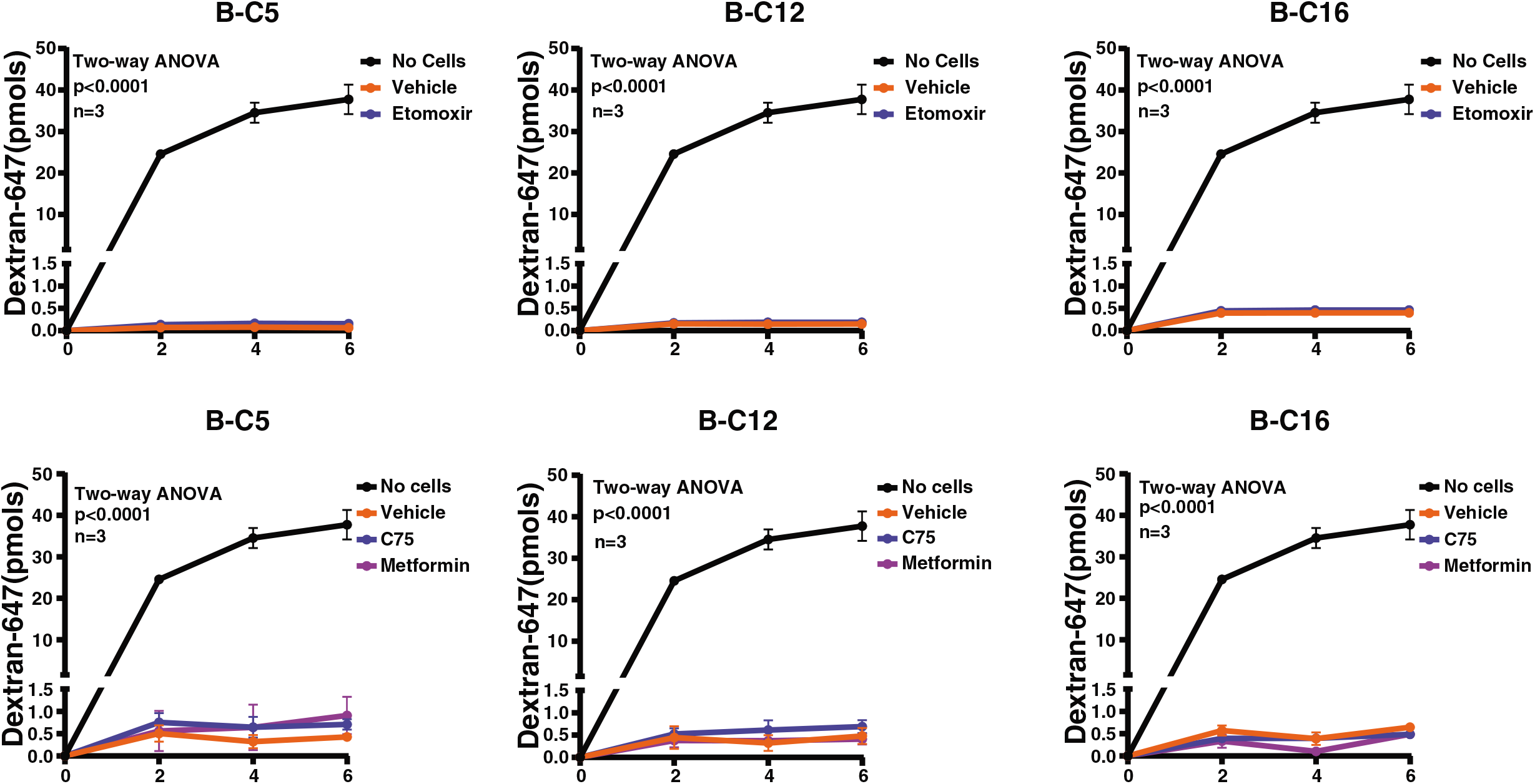
Quantification of Dextran-647 in the basal reservoir of AE monolayers from basal samples of FA-handling screens taken at 2, 4 and 6 hours. ****p < 0.0001. Experiments were done in triplicate (n=3 Transwell cultures per condition).

**Supplemental Figure 6.**
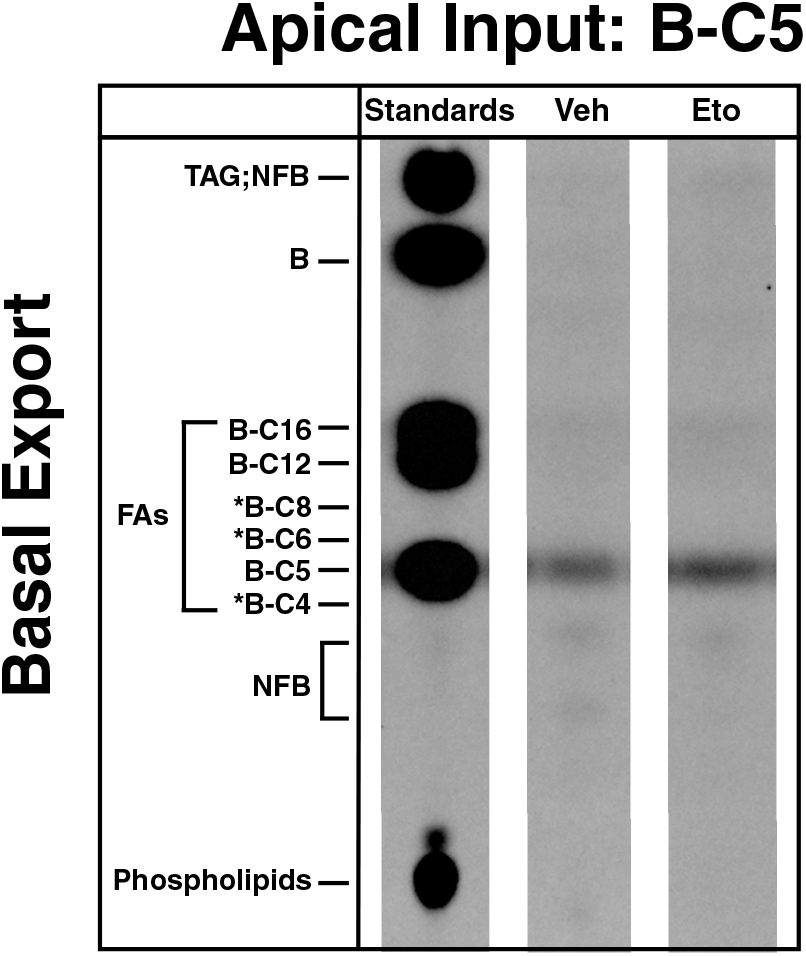
Thin layer chromatography was performed on the basal-reservoir media of B-C5 treated cultures treated with either vehicle or etomoxir. B-C5 and etomoxir concentrations were 20uM and 100uM respectively. A previous study^62^ using TLC to identify B-lipid species with the same solvents used in this study reported naturally fluorescent bands that do not correspond to B-lipid standards used and are labelled here as NFB. *Location of B-C4, B-C6 and B-C8 were inferred from the experiment in supplemental figure 4. Experiment was performed in triplicate (n = 3 AE transwell cultures).

## Acknowledgements

We would like to than the anonymous organ donors whose tissues were used in this study. We would like to thank the families of the organ donors as well as the organ procurement organization, HonorBridge (formerly Carolina Donor Services), for providing the tissue samples used in this study. We would like to thank the UNC Advanced Analytics Core for FACS and RNA-sequencing. Schematics were created with BioRender.com. This study was funded with NIH grants P30DK034987-35, R01DK115806, R01DK109559 and the Katherine Bullard Charitable Trust for Gastrointestinal Stem Cell and Regenerative Medicine Research. Scott Magness would like to disclose a financial interest in Altis Biosystems Inc., which licenses the ISC monolayer technology used in this study.

## References

1. Wang TY, Liu M, Portincasa P, Wang DQH. New insights into the molecular mechanism of intestinal fatty acid absorption. European Journal of Clinical Investigation 2013;43:1203–1223.

2. Ko CW, Qu J, Black DD, Tso P. Regulation of intestinal lipid metabolism: current concepts and relevance to disease. Nature Reviews Gastroenterology and Hepatology 2020;17:169–183.

3. McDonald GB, Saunders DR, Weidman M, Fisher L. Portal venous transport of long-chain fatty acids absorped from rat intestine. American Journal of Physiology - Gastrointestinal and Liver Physiology 1980;2:239.

4. Mansbach CM, Dowell RF, Pritchett D. Portal transport of absorbed lipids in rats. American Journal of Physiology - Gastrointestinal and Liver Physiology 1991;261:G530–G538.

5. Mu H, Høy CE. The digestion of dietary triacylglycerols. Progress in Lipid Research 2004;43:105–133.

6. Batterink L, Yokum S, Stice E. Body mass correlates inversely with inhibitory control in response to food among adolescent girls: An fMRI study. NeuroImage 2010;52:1696–1703.

7. Donofry SD, Stillman CM, Erickson KI. A review of the relationship between eating behavior, obesity and functional brain network organization. Social Cognitive and Affective Neuroscience 2020;15:1157–1181.

8. Athyros VG, Doumas M, Imprialos KP, Stavropoulos K, Georgianou E, Katsimardou A, Karagiannis A. Diabetes and lipid metabolism. Hormones 2018;17:61–67.

9. Glovaci D, Fan W, Wong ND. Epidemiology of Diabetes Mellitus and Cardiovascular Disease. Current Cardiology Reports 2019;21.

10. Rijn JM van, Ardy RC, Kuloglu Z, et al. Intestinal Failure and Aberrant Lipid Metabolism in Patients With DGAT1 Deficiency. Gastroenterology 2018;155:130–143.e15.

11. Luglio HF. Genetic variation of fatty acid oxidation and obesity, a literature review. International Journal of Biomedical Science 2016;12:1–8.

12. Ko C-W, Qu J, Black DD, Tso P. Regulation of intestinal lipid metabolism: current concepts and relevance to disease. Nature Reviews Gastroenterology & Hepatology 2020;17:169–183.

13. McCann JR, Bihlmeyer NA, Roche K, et al. The Pediatric Obesity Microbiome and Metabolism Study (POMMS): Methods, Baseline Data, and Early Insights. Obesity 2021;29:569–578.

14. Jumpertz R, Le DS, Turnbaugh PJ, Trinidad C, Bogardus C, Gordon JI, Krakoff J. Energy-balance studies reveal associations between gut microbes, caloric load, and nutrient absorption in humans. The American Journal of Clinical Nutrition 2011;94:58–65.

15. Nagpal R, Wang S, Solberg Woods LC, Seshie O, Chung ST, Shively CA, Register TC, Craft S, McClain DA, Yadav H. Comparative microbiome signatures and short-chain fatty acids in mouse, rat, non-human primate, and human feces. Frontiers in Microbiology 2018;9:2897.

16. Ridaura VK, Faith JJ, Rey FE, et al. Gut microbiota from twins discordant for obesity modulate metabolism in mice. Science 2013;341.

17. Trotter PJ, Ho SY, Storch J. Fatty acid uptake by Caco-2 human intestinal cells. Journal of Lipid Research 1996;37:336–346.

18. Bens M, Bogdanova A, Cluzeaud F, Miquerol L, Kerneis S, Kraehenbuhl JP, Kahn A, Pringault E, Vandewalle A. Transimmortalized mouse intestinal cells (m-IC(cl2)) that maintain a crypt phenotype. American Journal of Physiology - Cell Physiology 1996;270:C1666–C1674.

19. Pandrea IV, Carrière V, Barbat A, Cambier D, Dussaulx E, Lesuffleur T, Rousset M, Zweibaum A. Postmitotic differentiation of colon carcinoma Caco-2 cells does not prevent reentry in the cell cycle and tumorigenicity. Experimental and Molecular Pathology 2000;69:37–45.

20. Sun H, Chow ECY, Liu S, Du Y, Pang KS. The Caco-2 cell monolayer: usefulness and limitations. Expert opinion on drug metabolism & toxicology 2008;4:395–411.

21. Rahman S, Ghiboub M, Donkers JM, Steeg E van de, Tol EAF van, Hakvoort TBM, Jonge WJ de. The Progress of Intestinal Epithelial Models from Cell Lines to Gut-On-Chip. International Journal of Molecular Sciences 2021;22:13472.

22. Schutgens F, Clevers H. Human Organoids: Tools for Understanding Biology and Treating Diseases. Annual Review of Pathology: Mechanisms of Disease 2020;15:211–234.

23. Sato T, Vries RG, Snippert HJ, Wetering M Van De, Barker N, Stange DE, Es JH Van, Abo A, Kujala P, Peters PJ, Clevers H. Single Lgr5 stem cells build crypt-villus structures in vitro without a mesenchymal niche. Nature 2009;459:262–265.

24. Co JY, Margalef-Català M, Li X, Mah AT, Kuo CJ, Monack DM, Amieva MR. Controlling Epithelial Polarity: A Human Enteroid Model for Host-Pathogen Interactions. Cell Reports 2019;26:2509–2520.e4.

25. Rumjanek FD, Simpson AJG. The incorporation and utilization of radiolabelled lipids by adult Schistosoma mansoni in vitro. Molecular and Biochemical Parasitology 1980;1:31–44.

26. Murphy JL, Jones A, Brookes S, Wootton SA. The gastrointestinal handling and metabolism of [1-13C]palmitic acid in healthy women. Lipids 1995;30:291–298.

27. Kalivianakis M, Verkade HJ, Stellaard F, Werf M Van Der, Elzinga H, Vonk RJ. The 13C-mixed triglyceride breath test in healthy adults: determinants of the 13CO2 response. European Journal of Clinical Investigation 1997;27:434–442.

28. Hofmann M, Eichenberger W. Radiolabelling Studies on the Lipid Metabolism in the Marine Brown Alga Dictyopteris membranacea. Plant Cell Physiol 1998;39:508–515.

29. Ecker J, Liebisch G. Application of stable isotopes to investigate the metabolism of fatty acids, glycerophospholipid and sphingolipid species. Progress in Lipid Research 2014;54:14–31.

30. Triebl A, Wenk MR. Analytical considerations of stable isotope labelling in lipidomics. Biomolecules 2018;8:151.

31. Twining CW, Taipale SJ, Ruess L, Bec A, Martin-Creuzburg D, Kainz MJ. Stable isotopes of fatty acids: Current and future perspectives for advancing trophic ecology. Philosophical Transactions of the Royal Society B: Biological Sciences 2020;375.

32. Maier O, Oberle V, Hoekstra D. Fluorescent lipid probes: some properties and applications (a review). Chemistry and Physics of Lipids 2002;116:3–18.

33. Johnson ID, Kang HC, Haugland RP. Fluorescent membrane probes incorporating dipyrrometheneboron difluoride fluorophores. Analytical Biochemistry 1991;198:228–237.

34. Fredrik Bergström †, Ilya Mikhalyov †‡, Peter Hägglöf §, Rüdiger Wortmann ┴, Tor Ny § and, Lennart B.-Å. Johansson* †. Dimers of Dipyrrometheneboron Difluoride (BODIPY) with Light Spectroscopic Applications in Chemistry and Biology. Journal of the American Chemical Society 2001;124:196–204.

35. Valm AM, Cohen S, Legant WR, Melunis J, Hershberg U, Wait E, Cohen AR, Davidson MW, Betzig E, Lippincott-Schwartz J. Applying systems-level spectral imaging and analysis to reveal the organelle interactome. Nature 2017;546:162–167.

36. Rambold AS, Cohen S, Lippincott-Schwartz J. Fatty Acid Trafficking in Starved Cells: Regulation by Lipid Droplet Lipolysis, Autophagy, and Mitochondrial Fusion Dynamics. Developmental Cell 2015;32:678–692.

37. Wang Y, DiSalvo M, Gunasekara DB, Dutton J, Proctor A, Lebhar MS, Williamson IA, Speer J, Howard RL, Smiddy NM, Bultman SJ, Sims CE, Magness ST, Allbritton NL. Self-renewing Monolayer of Primary Colonic or Rectal Epithelial Cells. Cellular and Molecular Gastroenterology and Hepatology 2017;4:165–182.e7.

38. Hinman SS, Wang Y, Kim R, Allbritton NL. In vitro generation of self-renewing human intestinal epithelia over planar and shaped collagen hydrogels. Nature Protocols 2020;16:352–382.

39. Lema I, Araújo JR, Rolhion N, Demignot S. Jejunum: The understudied meeting place of dietary lipids and the microbiota. Biochimie 2020;178:124–136.

40. Busslinger GA, Weusten B LA, Bogte A, Begthel H, Brosens LA, Clevers H. Human gastrointestinal epithelia of the esophagus, stomach, and duodenum resolved at single-cell resolution. Cell Reports 2021;34.

41. Wang Y, Song W, Wang J, Wang T, Xiong X, Qi Z, Fu W, Yang X, Chen Y-G. Single-cell transcriptome analysis reveals differential nutrient absorption functions in human intestine. Journal of Experimental Medicine 2020;217.

42. Elmentaite R, Kumasaka N, Roberts K, et al. Cells of the human intestinal tract mapped across space and time. Nature 2021;597:250–255.

43. Burclaff J, Bliton RJ, Breau KA, Ok MT, Gomez-Martinez I, Ranek JS, Bhatt AP, Purvis JE, Woosley JT, Magness ST. A proximal-to-distal survey of healthy adult human small intestine and colon epithelium by single-cell transcriptomics. bioRxiv 2021:2021.10.05.460818.

44. Traag VA, Waltman L, Eck NJ van. From Louvain to Leiden: guaranteeing well-connected communities. Scientific Reports 2019;9:1–12.

45. Triana S, Stanifer ML, Metz-Zumaran C, Shahraz M, Mukenhirn M, Kee C, Serger C, Koschny R, Ordoñez-Rueda D, Paulsen M, Benes V, Boulant S, Alexandrov T. Single-cell transcriptomics reveals immune response of intestinal cell types to viral infection. Molecular Systems Biology 2021;17:9833.

46. Tirosh I, Izar B, Prakadan SM, et al. Dissecting the multicellular ecosystem of metastatic melanoma by single-cell RNA-seq. Science 2016;352:189–196.

47. Barker N, Es JH Van, Kuipers J, Kujala P, Born M Van Den, Cozijnsen M, Haegebarth A, Korving J, Begthel H, Peters PJ, Clevers H. Identification of stem cells in small intestine and colon by marker gene Lgr5. Nature 2007;449:1003–1007.

48. Muñoz J, Stange DE, Schepers AG, et al. The Lgr5 intestinal stem cell signature: robust expression of proposed quiescent ‘+4’’ cell markers.’ The EMBO Journal 2012;31:3079.

49. Gracz AD, Ramalingam S, Magness ST. Sox9 expression marks a subset of CD24-expressing small intestine epithelial stem cells that form organoids in vitro. American Journal of Physiology - Gastrointestinal and Liver Physiology 2010;298:G590.

50. Moor AE, Harnik Y, Ben-Moshe S, Massasa EE, Rozenberg M, Eilam R, Bahar Halpern K, Itzkovitz S. Spatial Reconstruction of Single Enterocytes Uncovers Broad Zonation along the Intestinal Villus Axis. Cell 2018;175:1156–1167.e15.

51. Manno G La, Soldatov R, Zeisel A, et al. RNA velocity of single cells. Nature 2018 560:7719 2018;560:494–498.

52. Bergen V, Lange M, Peidli S, Wolf FA, Theis FJ. Generalizing RNA velocity to transient cell states through dynamical modeling. Nature Biotechnology 2020;38:1408–1414.

53. Speer JE, Gunasekara DB, Wang Y, Fallon JK, Attayek PJ, Smith PC, Sims CE, Allbritton NL. Molecular transport through primary human small intestinal monolayers by culture on a collagen scaffold with a gradient of chemical cross-linking. Journal of Biological Engineering 2019;13.

54. Demitrack ES, Samuelson LC. Notch regulation of gastrointestinal stem cells. The Journal of Physiology 2016;594:4791–4803.

55. Shroyer NF, Helmrath MA, Wang VY –C., Antalffy B, Henning SJ, Zoghbi HY. Intestine-Specific Ablation of Mouse atonal homolog 1 (Math1) Reveals a Role in Cellular Homeostasis. Gastroenterology 2007;132:2478–2488.

56. Srinivasan B, Kolli AR, Esch MB, Abaci HE, Shuler ML, Hickman JJ. TEER measurement techniques for in vitro barrier model systems. Journal of laboratory automation 2015;20:107.

57. Wang Y, Kim R, Hwang SHJ, Dutton J, Sims CE, Allbritton NL. Analysis of Interleukin 8 Secretion by a Stem-Cell-Derived Human-Intestinal-Epithelial-Monolayer Platform. Analytical Chemistry 2018;90:11523–11530.

58. Otis JP, Farber SA. Imaging vertebrate digestive function and lipid metabolism in vivo. Drug Discovery Today: Disease Models 2013;10.

59. Anderson JL, Carten JD, Farber SA. Using fluorescent lipids in live zebrafish larvae: from imaging whole animal physiology to subcellular lipid trafficking. Methods in cell biology 2016;133:165.

60. Semova I, Carten JD, Stombaugh J, MacKey LC, Knight R, Farber SA, Rawls JF. Microbiota regulate intestinal absorption and metabolism of fatty acids in the zebrafish. Cell Host and Microbe 2012;12:277–288.

61. Sæle Ø, Rød KEL, Quinlivan VH, Li S, Farber SA. A novel system to quantify intestinal lipid digestion and transport. Biochimica et Biophysica Acta - Molecular and Cell Biology of Lipids 2018;1863:948–957.

62. Carten JD, Bradford MK, Farber SA. Visualizing digestive organ morphology and function using differential fatty acid metabolism in live zebrafish. Developmental Biology 2011;360:276.

63. Furse S, Kroon AIPM De. Phosphatidylcholines functions beyond that of a membrane brick. Molecular Membrane Biology 2015;32:117–119.

64. Houten SM, Violante S, Ventura F V., Wanders RJA. The Biochemistry and Physiology of Mitochondrial Fatty Acid β-Oxidation and Its Genetic Disorders. Annual Review of Physiology 2016;78:23–44.

65. Khuwijitjaru P, Adachi S, Matsuno R. Solubility of Saturated Fatty Acids in Water at Elevated Temperatures Materials and Methods. Biosci Biotechnol Biochem 2002;66:1723–1726.

66. Thupari JN, Landree LE, Ronnett G V, Kuhajda FP. C75 increases peripheral energy utilization and fatty acid oxidation in diet-induced obesity. Proceedings of the National Academy of Sciences of the United States of America 2002;99:9498–9502.

67. Wang C, Liu F, Yuan Y, Wu J, Wang H, Zhang L, Hu P, Li Z, Li Q, Ye J. Metformin suppresses lipid accumulation in skeletal muscle by promoting fatty acid oxidation. Clinical Laboratory 2014;60:887–896.

68. Zabielski P, Chacinska M, Charkiewicz K, Baranowski M, Gorski J, Blachnio-Zabielska AU. Effect of metformin on bioactive lipid metabolism in insulin-resistant muscle. Journal of Endocrinology 2017;233:329–340.

69. Kuncewitch M, Yang WL, Jacob A, Khader A, Giangola M, Nicastro J, Coppa GF, Wang P. Inhibition of fatty acid synthase with C75 decreases organ injury after hemorrhagic shock. Surgery (United States) 2016;159:570–579.

70. Zachos NC, Kovbasnjuk O, Foulke-Abel J, In J, Blutt SE, Jonge HR de, Estes MK, Donowitz M. Human Enteroids/Colonoids and Intestinal Organoids Functionally Recapitulate Normal Intestinal Physiology and Pathophysiology. The Journal of Biological Chemistry 2016;291:3759.

71. Raab JR, Tulasi DY, Wager KE, Morowitz JM, Magness ST, Gracz AD. Quantitative classification of chromatin dynamics reveals regulators of intestinal stem cell differentiation. Development (Cambridge) 2020;147.

72. Kim T-H, Li F, Ferreiro-Neira I, Ho L-L, Luyten A, Nalapareddy K, Long H, Verzi M, Shivdasani RA. Broadly permissive intestinal chromatin underlies lateral inhibition and cell plasticity. Nature 2014;506:511.

73. Carlier H, Bezard J. Electron microscope autoradiographic study of intestinal absorption of decanoic and octanoic acids in the rat. Journal of Cell Biology 1975;65:383–397.

74. Ways PO, Parmentier CM, Kayden HJ, Jones JW, Saunders DR, Rubin CE. Studies on the Absorptive Defect for Triglyceride in Abetalipoproteinemia. The Journal of Clinical Investigation 1967;46:35–46.

75. Christensen LW, Kuhre RE, Janus C, Svendsen B, Holst JJ. Vascular, but not luminal, activation of FFAR1 (GPR40) stimulates GLP-1 secretion from isolated perfused rat small intestine. Physiological Reports 2015;3.

76. Kimura I, Ichimura A, Ohue-Kitano R, Igarashi M. Free fatty acid receptors in health and disease. Physiological Reviews 2020;100:171–210.

77. Bahne E, Sun EWL, Young RL, et al. Metformin-induced glucagon-like peptide-1 secretion contributes to the actions of metformin in type 2 diabetes. JCI insight 2018;3.

78. Mulherin AJ, Oh AH, Kim H, Grieco A, Lauffer LM, Brubaker PL. Mechanisms Underlying Metformin-Induced Secretion of Glucagon-Like Peptide-1 from the Intestinal L Cell. Endocrinology 2011;152:4610–4619.

79. Kowalczyk MS, Tirosh I, Heckl D, Rao TN, Dixit A, Haas BJ, Schneider RK, Wagers AJ, Ebert BL, Regev A. Single-cell RNA-seq reveals changes in cell cycle and differentiation programs upon aging of hematopoietic stem cells. Genome Research 2015;25:1860.

80. Wolf FA, Angerer P, Theis FJ. SCANPY: Large-scale single-cell gene expression data analysis. Genome Biology 2018;19:1–5.

81. Traag VA, Waltman L, Eck NJ van. From Louvain to Leiden: guaranteeing well-connected communities. Scientific Reports 2019;9:1–12.

82. Wolf FA, Hamey FK, Plass M, Solana J, Dahlin JS, Göttgens B, Rajewsky N, Simon L, Theis FJ. PAGA: graph abstraction reconciles clustering with trajectory inference through a topology preserving map of single cells. Genome Biology 2019;20:1–9.

83. Fischer DS, Hölzlwimmer FR, Eraslan G, Heumos L. Fast and scalable differential expression analysis on single-cell RNA-seq data. 2020. Available at: https://github.com/theislab/diffxpy.

84. Schroeder A, Mueller O, Stocker S, Salowsky R, Leiber M, Gassmann M, Lightfoot S, Menzel W, Granzow M, Ragg T. The RIN: An RNA integrity number for assigning integrity values to RNA measurements. BMC Molecular Biology 2006;7.

85. Robinson MD, Oshlack A. A scaling normalization method for differential expression analysis of RNA-seq data. Genome Biology 2010;11.

86. Bray NL, Pimentel H, Melsted P, Pachter L. Near-optimal probabilistic RNA-seq quantification. Nature Biotechnology 2016;34.

87. Sündermann A, Eggers LF, Schwudke D. Liquid Extraction: Bligh and Dyer. Encyclopedia of Lipidomics 2016:1–4.

